# Stepwise evolution and convergent recombination underlie the global dissemination of carbapenemase-producing *Escherichia coli*

**DOI:** 10.1101/446195

**Authors:** Rafael Patiño-Navarrete, Isabelle Rosinski-Chupin, Nicolas Cabanel, Lauraine Gauthier, Julie Takissian, Jean-Yves Madec, Monzer Hamze, Remy A. Bonnin, Thierry Naas, Philippe Glaser

## Abstract

Carbapenem-resistant *Enterobacteriaceae* are considered by WHO as “critical” priority pathogens for which novel antibiotics are urgently needed. The dissemination of carbapenemase-producing *Escherichia coli* (CP-*Ec*) in the community is a major public health concern. However, the global molecular epidemiology of CP-*Ec* isolates, as well as the genetic bases for the emergence and global dissemination of specific lineages, remain largely unknown. Here, by combining a thorough genomic and evolutionary analysis of *Ec* ST410 isolates with a broad analysis of 12,584 *E. coli and Shigella* genomes, we showed that the fixation of carbapenemase genes depends largely on a combination of mutations in *ftsI* encoding the penicillin binding protein 3 and in the porin genes *ompC* and *ompF*. Mutated *ftsI* genes and a specific *ompC* allele inducing reduced susceptibility to diverse β-lactams spread across the species by recombination. The selection of CP-*Ec* lineages able to disseminate is more complex than the mere acquisition of carbapenemase genes.

## Introduction

Antibiotic resistance is one of the most urgent public health concerns. The increasing rate in antimicrobial resistances worldwide suggests a bleak outlook in terms of morbidity, mortality and economic loss^1^. Carbapenems are one of the last resort antibiotics used to treat infections caused by multidrug-resistant (MDR) Gram-negative bacteria^2^. Dissemination of carbapenem-resistant *Enterobacteriaceae* (CRE) threatens the efficacy of current treatment options. Carbapenem-resistance may result from a combination of mutations leading to reduced permeability (e.g. porin deficiency) and overexpression of an extended-spectrum β-lactamase (ESBL) or a cephalosporinase that show a weak activity against carbapenems^3^. However, the main resistance mechanism is the acquisition of a carbapenemase gene^4^. The major carbapenemases encountered in *Enterobacteriaceae* belong to Ambler class A (KPC-type), class B (metallo-β-lactamases IMP, VIM- and NDM-types) or class D (OXA-48-like enzymes)^5^. As these carbapenemases are now frequently encountered in *Escherichia coli*, carbapenemase-producing *E. coli* (CP-*Ec*) might follow the same expansion and dissemination in hospitals and the community as the one observed for CTX-M-type ESBL-producing *E. coli* isolates^6,7^. A scenario feared by public health authorities. This is especially worrisome as these isolates are usually resistant to multiple antibiotics.

The epidemiology of CP-*Ec* is complex with geographical diversity in terms of carbapenemase genes and of dominant lineages^4^. Most studies performed at national or hospital levels point to a broad diversity of isolates as defined by multilocus sequence typing (MLST), with some isolates belonging to a few dominant sequence types (STs) like ST38, clonal complex (CC) 10 (ST10, ST167, ST617), ST101, ST131 and ST410 that express different carbapenemases^4,8–14^. However, their prevalence varies significantly worldwide. Analysis of CP-*Ec* strains isolated in 16 countries between 2008 and 2013 revealed that 36% belonged to the pandemic ST131, which has driven the global spread of CTX-M-15 ESBL in *E. coli*^11^. Similarly, a survey of CRE strains in China showed that ST131 represented 34% of the isolates and ST167, 17%^14^. But only a single ST131 isolate out of 140 CP-*Ec* was identified by the French National Reference Centre (Fr-NRC) between 2012-2013^8^. Recently, the phylogenetic analysis of a Danish collection of ST410 isolates combined to an international set of isolates revealed a globally disseminated *Ec* clone carrying *bla*_OXA-181_ on a IncX3 plasmid. This lineage was predicted by a Bayesian analysis to have acquired *bla*_OXA-181_ around 2003 and subsequently *bla*_NDM—5_ around 2014^13^.

Despite public health implications, factors contributing to the emergence and the dissemination of CP-*Ec* lineages have not been explored. Here, by using an in-depth evolutionary and functional analysis of *Ec* ST410 and by extending it to the whole *E. coli* species, we show that acquisition of carbapenemase genes followed different evolutionary trajectories. In most STs it occurred preferentially in specific disseminated lineages mutated in *ftsI* encoding penicillin-binding protein 3 (PBP3) and/or in the *ompC* and *ompF* porin genes. We also show that these mutations lead to a reduced susceptibility to some β-lactams including ertapenem. On the other hand, we did not identify these mutations among ST131 isolates. These new data on evolution of CP-*Ec* allow us to propose a model for their selection and dissemination.

## Results

### Most CP-*Ec* ST410 isolates received by the French NRC belong to a single lineage

In order to determine the genetic bases for the dissemination of CP-*Ec* lineages, we first analysed ST410 CP-*Ec* isolates, which show a high prevalence among isolates collected by the Fr-NRC^8^. We sequenced the genomes of 54 CP-Ec isolates, 50 collected by the Fr-NRC (including 22 from patients repatriated from 15 different countries) and four from Lebanon, and three non-CP isolates of animal origin (Supplementary Table 1). We reconstructed their phylogeny together with 149 *Ec* ST410 genome sequences retrieved from public databases (Supplementary table 2). We filtered for redundancy in this collection by removing 50 clonal isolates differing by less than seven SNPs in the core genome^15^ and keeping the isolate with the largest number of antibiotic resistance genes (ARG). The phylogeny is in agreement with the recent analysis of CP-*Ec* ST410 from a Danish collection^13^, with a major fluoroquinolone resistant clade (FQR-clade) gathering the majority of the non-redundant (nr) isolates (133 out of 155) and of the nr-isolates carrying carbapenemase genes (62 out of 63). Within the FQR-clade 77% of the isolates expressed CTX-M-type ESBLs (Fig. 1). 36 out of the 40 *bla*_OXA-181_ carrying isolates formed a single subclade (the OXA-181 subclade) which corresponds to the previously described clade B4/H24RxC^13^. The 24 CP-*Ec* isolates not belonging to the OXA-181 subclade express different carbapenemases from the OXA-48, KPC, VIM and NDM families.

**Fig. 1.**
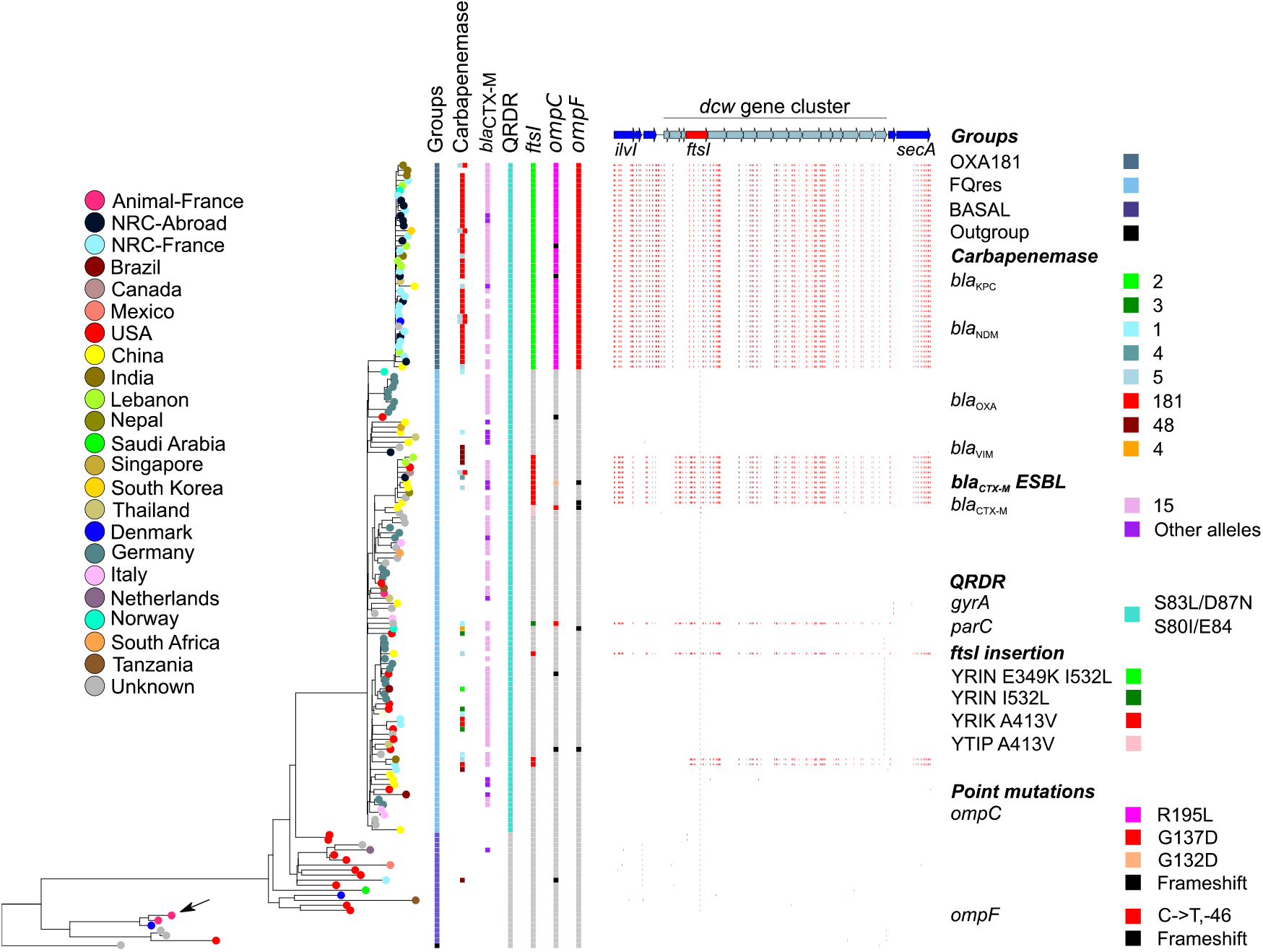
Core genome phylogeny and genomic features of *E. coli* ST410 isolates. ML phylogeny of 155 *Ec* ST410 nr-genomes built with RAxML^16^ based on the 3,419,955 bp core and recombination-free alignment of 5,378 SNPs. The *Ec* ST88 isolate 789 (CP010315.1) was used as an outgroup. Isolates (branch tips) are colour-coded according to the geographical origin as indicated in the figure key (left). Genomic features are indicated as indicated in the figure key (right) from left to right: groups according to the phylogeny, including the FQR clade and the OXA-181 subclade; carbapenemases; CTX-M ESBL; mutations in *gyrA* and *parC* QRDR region (FQ resistance); mutations in *ftsI*, *ompC,* and *ompF*. SNPs in the *dcw* cluster compared to the *Ec* ST410 non-recombined strain ANSES30599 (black arrow) are indicated by small vertical red bars. Upper part, genetic map of the *dcw* locus, genes are indicated by arrows, *ftsI* in red. NRC stands for National Reference Centre.

To accurately analyse the evolution of the OXA-181 subclade, we sequenced to completion a representative isolate of this clade (*Ec*-MAD). *Ec*-MAD carries three plasmids and 16 ARGs targeting seven classes of antibiotics (Supplementary Table 3). Indeed, antibiotic susceptibility testing showed that it is resistant to most antibiotics tested, remaining susceptible only to imipenem, meropenem, doripenem, amikacin, azithromycin, chloramphenicol, tigecycline, and colistin and intermediate to mecillinam, ertapenem, kanamycin and gentamicin (Supplementary Table 4). Comparison of the ARG content among ST410 *Ec* isolates revealed an increase in the median number of ARG between the basal isolates (n=4), the FQR-clade (n=9) and the OXA-181 subclade (n=16) (Supplementary Fig. 1).

### Gain of specific *ftsI* alleles by recombination is a hallmark of *Ec* ST410 carbapenemase-producing strains

Our phylogenetic analysis provided further evidence of a worldwide dissemination of the OXA-181 subclade^13^. Therefore, we searched for polymorphisms that, in addition to the acquisition of ARGs, have contributed to the expansion of this lineage. To this end, we systematically analysed mutations occurring in the branch leading to its most recent common ancestor (MRCA). Besides 97 mutations in non-recombined regions, we also identified 1648 SNPs in regions predicted as recombined by using Gubbins^17^ (Supplementary Table 5). 92,4 % occurred in a 124-kb DNA region between *yaaU* and *erpA* (Supplementary Fig. 2). In contrast, this recombined region was almost identical to sequences found in four ST167 and eight ST617 isolates from CC 10. Strikingly, all but one of these isolates expressed a carbapenemase gene. Furthermore, analysis of ST410 CP-*Ec* isolates outside the OXA-181 subclade revealed four additional recombination events overlapping the 124-kb recombined region identified in the OXA-181 subclade (Supplementary Fig. 2, Fig. 1). These recombination events affected a subclade of ten nr-isolates of different geographical origins including five CP isolates carrying different carbapenemase genes; two closely related CP-*Ec* isolates, one from India (*bla*NDM-5) and one from the Fr-NRC (*bla*_OXA-181_); and isolated CP-*Ec* isolates (Fig. 1). The 16.5-kb region shared by the five recombined regions encompassed the *dcw* (division and cell wall) locus from *ftsI* to *secM* (Fig. 1). It encodes major functions in cell wall synthesis and cell division, including *ftsI* encoding PBP3, a target of diverse β-lactams^18^. Overall, 75% (47/63) of the nr CP-*Ec* ST410 isolates had recombined in the *dcw* region (Fig. 1).

197 SNPs including 16 non-synonymous (NS) mutations differentiated the common 16.5-kb region in the OXA-181 subclade from other *Ec* ST410 isolates (Supplementary Table 5). Among differences, we identified insertions of four codons (YRIN in two cases and YRIK in three, Fig. 2a) occurring at a same position, between P333 and Y334 of FtsI. These insertions resulting from a four codons duplication (YRIN) and from a subsequent mutation (YRIK) were first described in NDM-producing *E. coli* isolates from different STs^19^. Additional NS SNPs were identified in the *ftsI* gene: E349K and I532L in association with the YRIN insertion and A413V with the YRIK insertion. The YRIK insertion in PBP3 was previously shown to confer reduced susceptibility to different β-lactams including ampicillin, cefepime and aztreonam but not to carbapenems^19^.

**Fig. 2.**
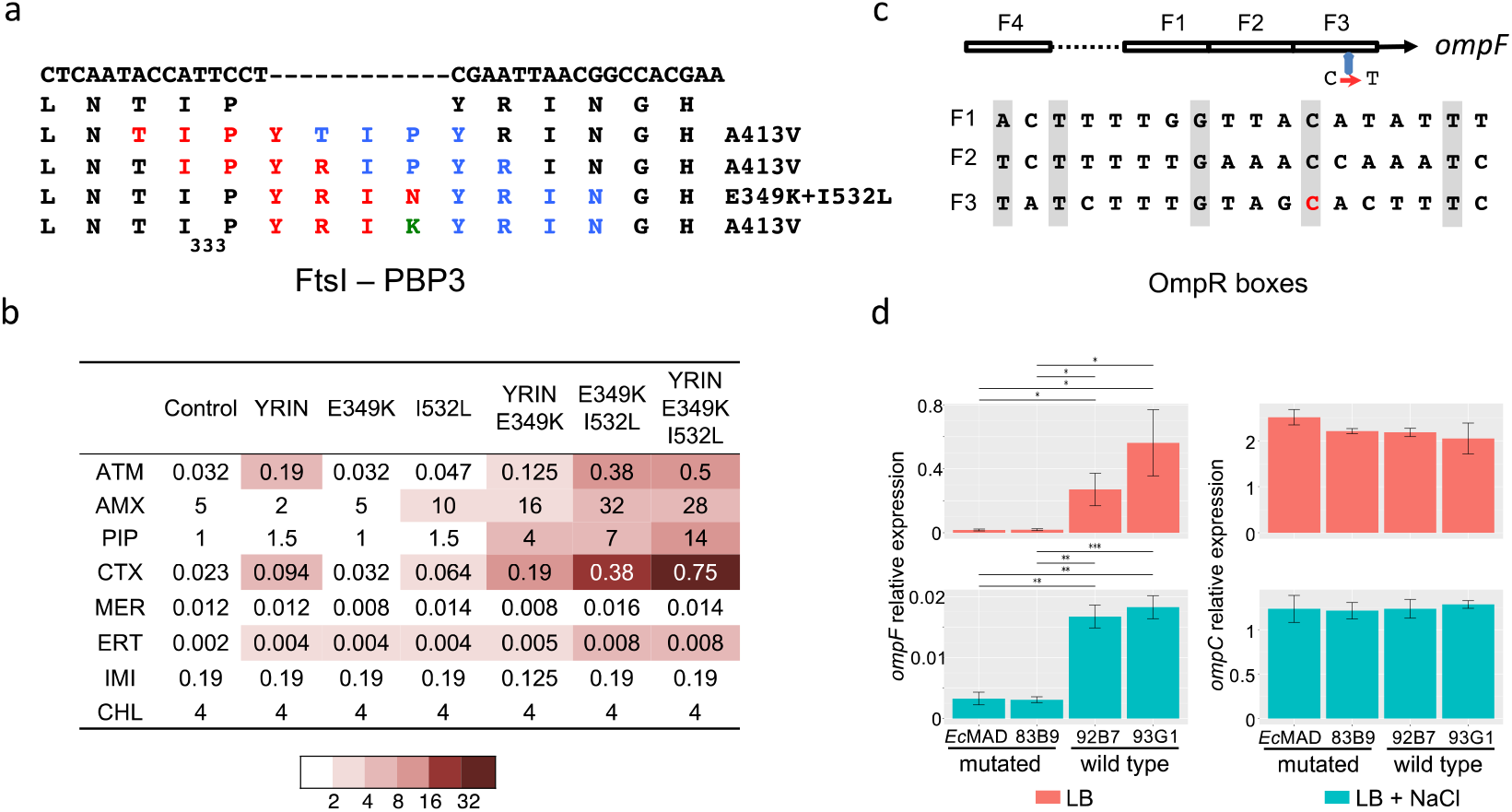
Functional analyses of *ftsI* and *ompF* mutations occurring in the OXA-181 *Ec* ST410 subclade. **a.** Mutations identified in *ftsI*. The four different insertions following proline 333 resulted in the duplication of the four codons shown in red and blue. The YRIK insertion derived from YRIN by a N to K AA change (in green). The first and second lines represent the WT nucleotide and AA sequences respectively; on the right, AA substitutions associated with each duplication. **b.** Antibiotic susceptibility testing performed by Etest of MG1655 derivatives mutated in *ftsI*. Abbreviation: ATM, Aztreonam; AMX, Amoxicillin; PIP, Piperacillin; CTX, Cefotaxime; MER, Meropenem; ERT, Ertapenem; IMI, Imipenem; CHL, Chloramphenicol. **c.** Schematic representation of the four OmpR binding sites in the *ompF* regulatory region and mutation of the conserved cytosine (C=>T) in the F3 OmpR binding site in red. **d.** Expression of *ompF* and *ompC* genes in two strains from the OXA-181 subclade (*Ec*MAD and 83B9, mutated) or from the FQR clade (92B7 and 93G1, WT) grown in LB medium and in LB medium supplemented with 0.3M NaCl. Bars represent confidence intervals; *, P < 0.05; **, P < 0.01; ***, P < 0.001.

### Mutations in the porin genes *ompC* and *ompF* are predicted to also have contributed to the selection of the ST410 OXA-181 subclade

To identify additional polymorphisms that might have contributed to the dissemination of the *Ec* ST410 OXA-181 subclade, we analysed the potential effect of non-synonymous mutations in the branch leading to its MRCA by using the SIFT algorithm^20^. We identified 34 NS mutations with a predicted functional effect (nine in the recombined region) (Supplementary Table 6). Eight of these mutations affected genes from the class “transporter” including the multidrug efflux transporter components *emrD* and *emrK* and five from the class “cell envelope”. These mutations might have been selected in relation to modifications in antibiotic susceptibility.

Among mutations affecting functions related to the cell envelope, one was the *ftsI* mutation I532L; another affected the porin gene *ompC* at a conserved arginine residue in the L4 loop (R195L, OmpC MG1655 numbering), one of the gateways for carbapenems (Fig. 3a)^21^. Arg_195_ is exposed, at the vestibule of the pore lumen and is conserved in OmpF^22^. Therefore, its replacement by a leucine, a non-polar AA, might affect permeation of β-lactams into the periplasm as we confirmed experimentally (see below). While we did not detect mutations in *ompF* coding sequence in the OXA-181 subclade, we identified a mutation in *ompF* regulatory region. This mutation replaces a conserved cytosine to a thymine residue in the proximal (F3) OmpR binding site. OmpR is a transcriptional activator of *ompF* and *ompC* expression and this mutation is predicted to affect *ompF* expression (Fig. 2c)^23^.

**Fig 3.**
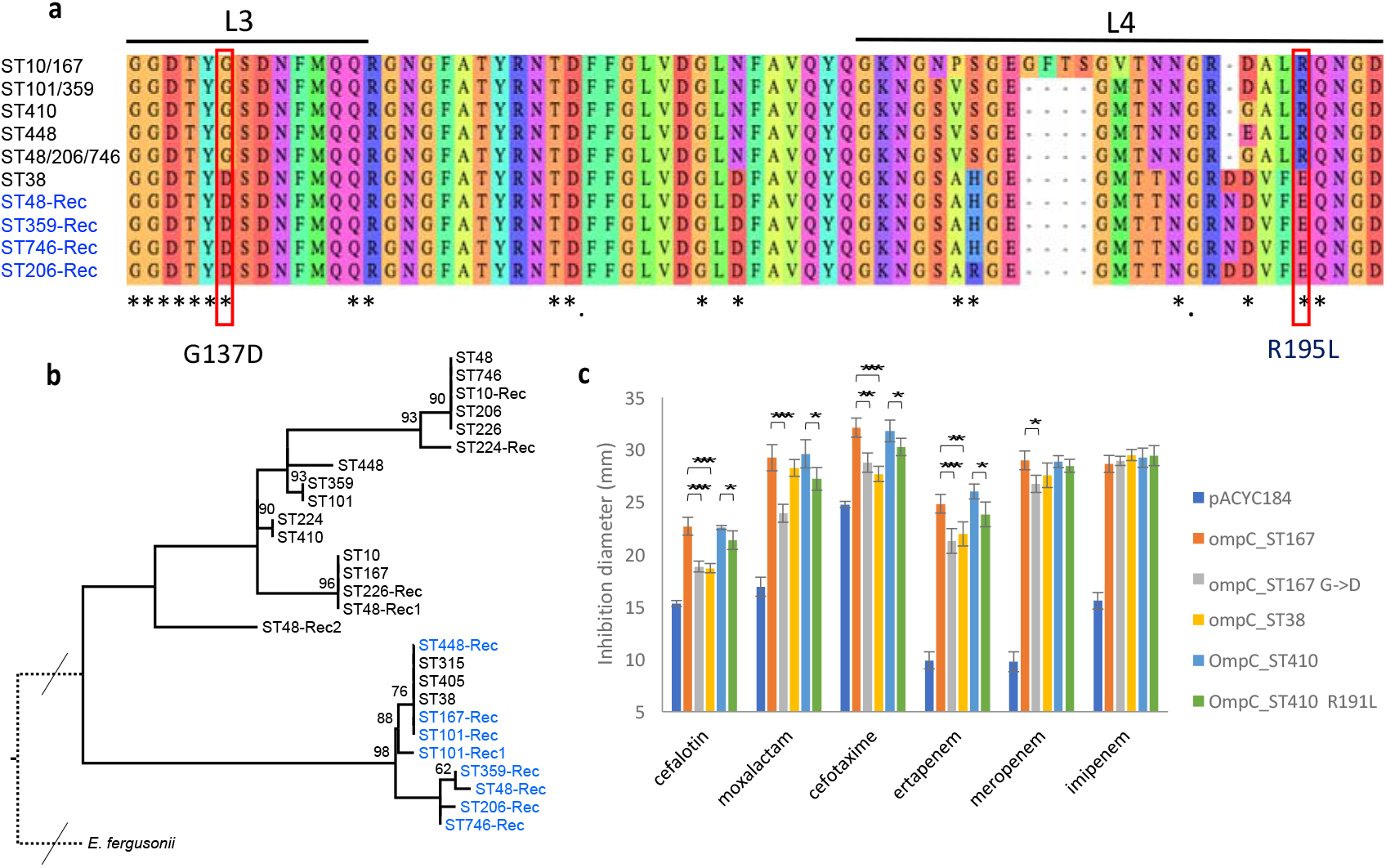
Mutations and recombination in the *ompC* gene. **a**. Alignment of OmpC L3L4 region from ST in which mutations or recombination events were detected. External loops L3 and L4 are indicated by lines above the sequences, and those positions predicted to be exposed to the pore lumen in *E. coli* MG1655 (ST10) by asterisks^22^. Mutation R195L and G137D associated with the gain of carbapenemase genes are highlighted by red rectangles. Numbering is according to MG1655 OmpC protein. **b.** Maximum likelihood phylogenetic reconstruction of representative OmpC sequences. OmpC sequences are labelled according to the ST of origin. OmpC sequences tagged with “-Rec” in blue were acquired by recombination in their respective STs; independent recombination events with different *ompC* alleles within a single ST are numbered. Bootstrap values > 60 are indicated. **c.** Antibiotic susceptibility testing (diameters of inhibition) of W3110 Δ*ompC* Δ*ompF* pOXA-232 strain complemented by different alleles of the *ompC* gene cloned in the medium copy number pACYC184^24^ according to the figure key. The empty vector was used as control. Bars represent standard deviations; *, P < 0.05; **, P < 0.01; ***, P < 0.001.

### Recombination at the *dcw* cluster and mutations in the porin genes *ompC* and *ompF* are frequently associated with the acquisition of a carbapenemase gene

In addition to the YRIN and YRIK insertions, two other four-AA insertions were also recently reported at the same position in FtsI, YRIP and YTIP. These insertions result from duplications starting two and three codons upstream the YRIN duplication respectively (Fig. 3a)^25^. To determine whether the association between the acquisition of a carbapenemase gene and a mutated PBP3 characterized by a four AA insertion is specific to ST410 isolates or if is also observed in other *E. coli* lineages, we analysed for these two features *E. coli* and *Shigella* genomes from the NCBI database. Indeed, despite biases of sequenced genomes deposited in public databases, they originated from different parts of the world and were analysed for different purposes and cover the diversity of the species. None of the *Shigella* isolates encoded a carbapenemase gene or carried an insertion in PBP3. 487 *E. coli* isolates (4.4%) encoded a carbapenemase gene and 248 (2.3%) carried a four AA insertion in PBP3: 163 YRIN, 49 YRIK, 3 YTIP and 33 YRIP. After removing redundancy for almost identical isolates of the same origin, 80% (146 out of 182) of nr-isolates mutated in *ftsI* were CP-*Ec* (Supplementary Table 7). All the 123 nr-isolates showing the YRIN insertion were also mutated at position 532 (I/L) and 112 at position 349 (E/K). On the other hand, all YRIK, YTIP and YRIP insertions were associated with a same secondary mutation A413V, suggesting this AA change was selected together with the four AA insertion either to reduce the fitness cost of the AA insertion or to reduce the susceptibility to antibiotics targeting PBP3. Globally, these data reveal at the species level, a strong link between these combinations of mutations in PBP3 and the acquisition of a carbapenemase gene. In addition to ST410, *ftsI* was mutated in the majority of the CP-*Ec* nr-isolates from ST101 (100%, N=23), ST167 (91%, N=49) and ST405 (81%, N=13) (Supplementary table 7). The case of the *Ec* ST167 isolates was particularly striking. We detected, within this single ST, after including 75 nr-isolates from Enterobase, 13 events of recombination leading to the replacement of the endogenous PBP3 allele by an allele with the YRIN (n=11) or the YRIK (n=2) insertion. Eleven recombination events affected at least one CP isolate, including a sub-clade of 35 nr-isolates expressing carbapenemases from seven different types. Only five out of the 54 *Ec* ST167 CP isolates did not undergo recombination at *ftsI* (Supplementary Fig. 3).

We next reconstructed the phylogeny of the 25 STs with at least 3 CP-*Ec* isolates and analysed mutations in *ompC* and *ompF* occurring during the evolution of these STs (Supplementary table 7). We focused on mutations inactivating *ompF* or decreasing its expression by affecting OmpR binding sites in the promoter region, as observed in the ST410 OXA-181 subclade. We also looked at mutations inactivating *ompC* or predicted to modify the porin permeability to β-lactams by decreasing the charge of AA located in the pore lumen^21^. 117 CP-*Ec* nr-isolates (41%) out of 286 from the NCBI were mutated in *ompF* compared to only 138 (8%) out of the 1659 non CP-Ec nr-isolates. This shows that these *ompF* alterations are associated with the acquisition of a carbapenemase gene. In 89 CP-*Ec* nr-isolates (31%), OmpC was modified but in only three CP-*Ec* (1%) isolates it was inactivated. In non CP-*Ec* isolates, OmpC was modified in only 44 (3%) nr-isolates and inactivated or missing in 39 (2%) (supplementary table 7). Therefore, OmpC modifications, but not its inactivation, are associated with the acquisition of a carbapenemase gene. This might be due to the high fitness cost of OmpC loss^26^. In addition to the R195L mutation in the OXA-181 ST410 subclade we identified two positions in the constriction loop L3 of OmpC^22^ independently mutated in different isolates. The G137D replacement was identified in a ST361 lineage enriched in CP-*Ec* isolates (Supplementary Fig. 4) and in four independent CP-*Ec* isolates from ST410, ST448 and ST617 (Fig. 1 and Supplementary Fig. 4), and the G132D in a carbapenemase resistant isolate belonging to a ST410 lineage mutated in *ftsI* and in two ST405 isolates (Fig. 1 and Supplementary Fig. 5). However, the most frequent *ompC* modification associated with CP-*Ec* isolates was the replacement of the original allele by alleles originating from phylogroup D strains through recombination (Fig. 3b). Indeed, we observed 16 independent recombination events, notably in the broadly distributed ST167 subcluster with a 22.7 kb recombined region from ST38 (Supplementary Fig. 6). Strikingly, OmpC proteins from phylogroup D isolates differ from other *E. coli* OmpC proteins at the two aforementioned residues G137 and R195 by negatively charged residues, D and E respectively (Fig. 3). In addition to ST38^8,9^ four other STs from phylogroup D: ST354, ST405, ST457 and ST648 contained CP-*Ec* isolates (Supplementary Fig. 5). These observations point to an association between this *ompC* allele and the acquisition of a carbapenemase gene.

Alike for *ompC* gene, we were able to trace recombination events leading to mutations in *ftsI* by analysing polymorphisms at the *dcw* locus considering genome-based phylogeny (Fig. 4 and supplementary Fig. 6). The identification of mutated genes with no signs of recombination (i.e. the absence of additional SNPs in *ftsI* and in the *dcw* gene cluster compared to basal strains) showed that YRIN, YRIP and YTIP insertions occurred first in one strain of the ST101, ST156 (Supplementary Fig. 3) and ST410 lineages (Fig. 1), respectively. The *ftsI* region with an YRIK insertion is almost identical to the ST101 *ftsI* region with YRIN insertion and therefore likely derived from it. The four *ftsI* variants were subsequently transferred to other lineages by lateral gene transfer (LGT). For instance, a 29.5 kb region recombined first from ST101 to ST167 and thereafter a 124-kb region from ST167 recombined into the MRCA of the ST410 OXA-181 subclade (Supplementary Fig. 6). Similarly, a 65-kb region with the YRIP insertion in *ftsI* from an ST156 *Ec* strain was introduced by homologous recombination in the MRCA of a clade of NDM-9 expressing ST224 *Ec* isolates. The shortest recombination event, detected in a ST167 CP-*Ec* strain (WCHEC16) contains only the *ftsI* gene (Supplementary Fig. 3). In total, we detected 52 independent recombination events involving a mutated *ftsI* allele scattered in all *E. coli* phylogroups except the B2 strains. Indeed, in *Ec* ST131 from the B2 phylogroup, despite the large number of CP-*Ec* isolates (n=49 nr-isolates), no isolate was mutated in *ftsI* (Supplementary Fig. 7).

**Fig. 4.**
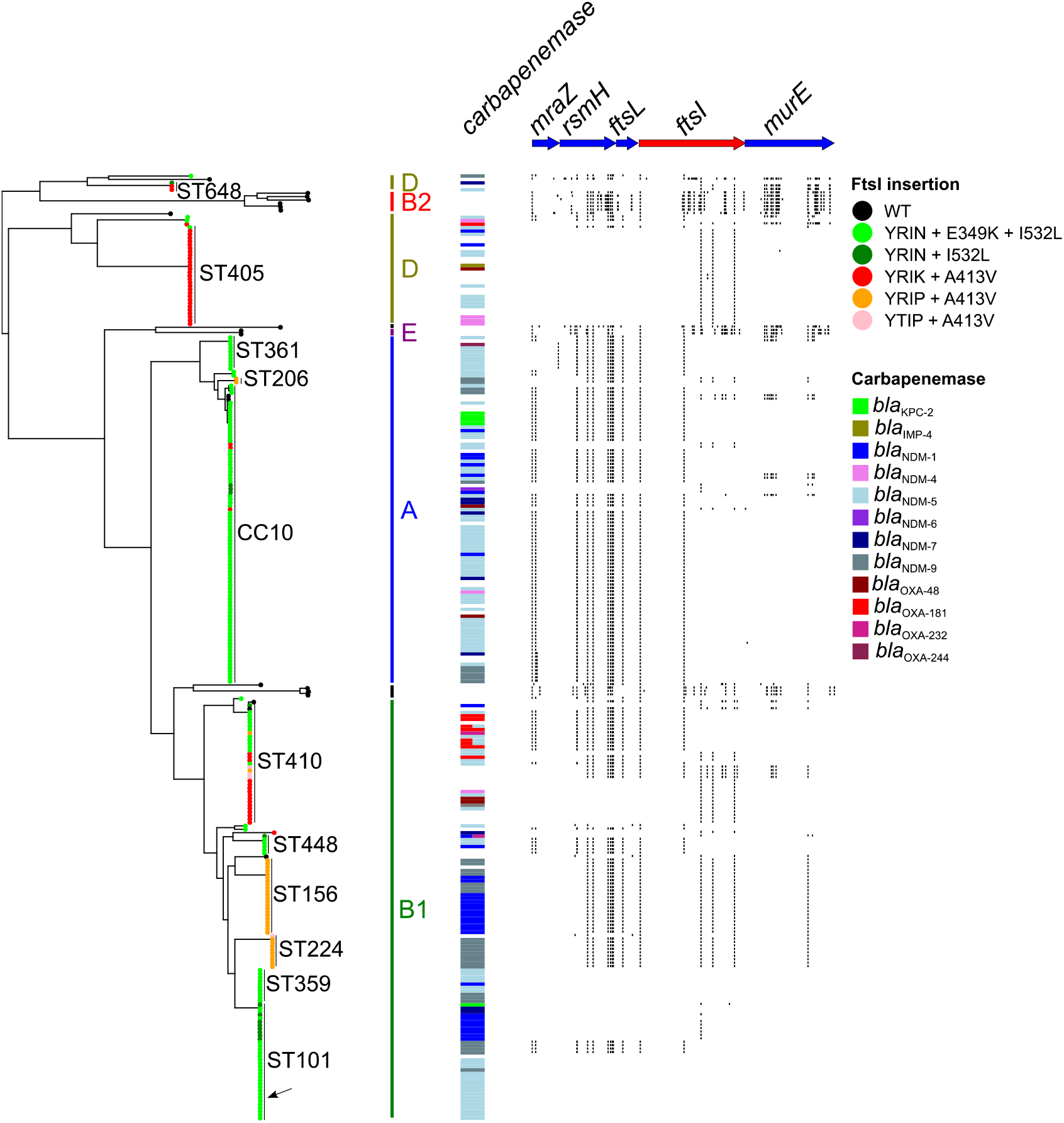
Phylogenetic distribution of *E. coli* isolates mutated in *ftsI*. The genome based phylogeny of *Ec* isolates mutated in *ftsI* and of *E. coli* reference strains^27^ was estimated from core non-recombinant SNP. Mutations in the *ftsI* gene are indicated at the tip of the branch as defined in the figure keys. Major STs are indicated, as well as phylogroups, S standing for *Shigella*. Carbapenemase genes are colour-coded as defined in the figure keys on the right. SNPs in the *mraZ* – *murE* region compared to the *Ec* ST101 strain CREC-591 (CP024821) shown by a black arrow are indicated by small vertical black bars. An ST101 isolate was chosen as reference, as insertion of YRIN in *ftsI* occurred among this ST. Conservation of SNP patterns between isolates showing the same mutation in *ftsI* across the *E. coli* species revealed the exchange of these alleles by recombination.

**Fig. 5.**
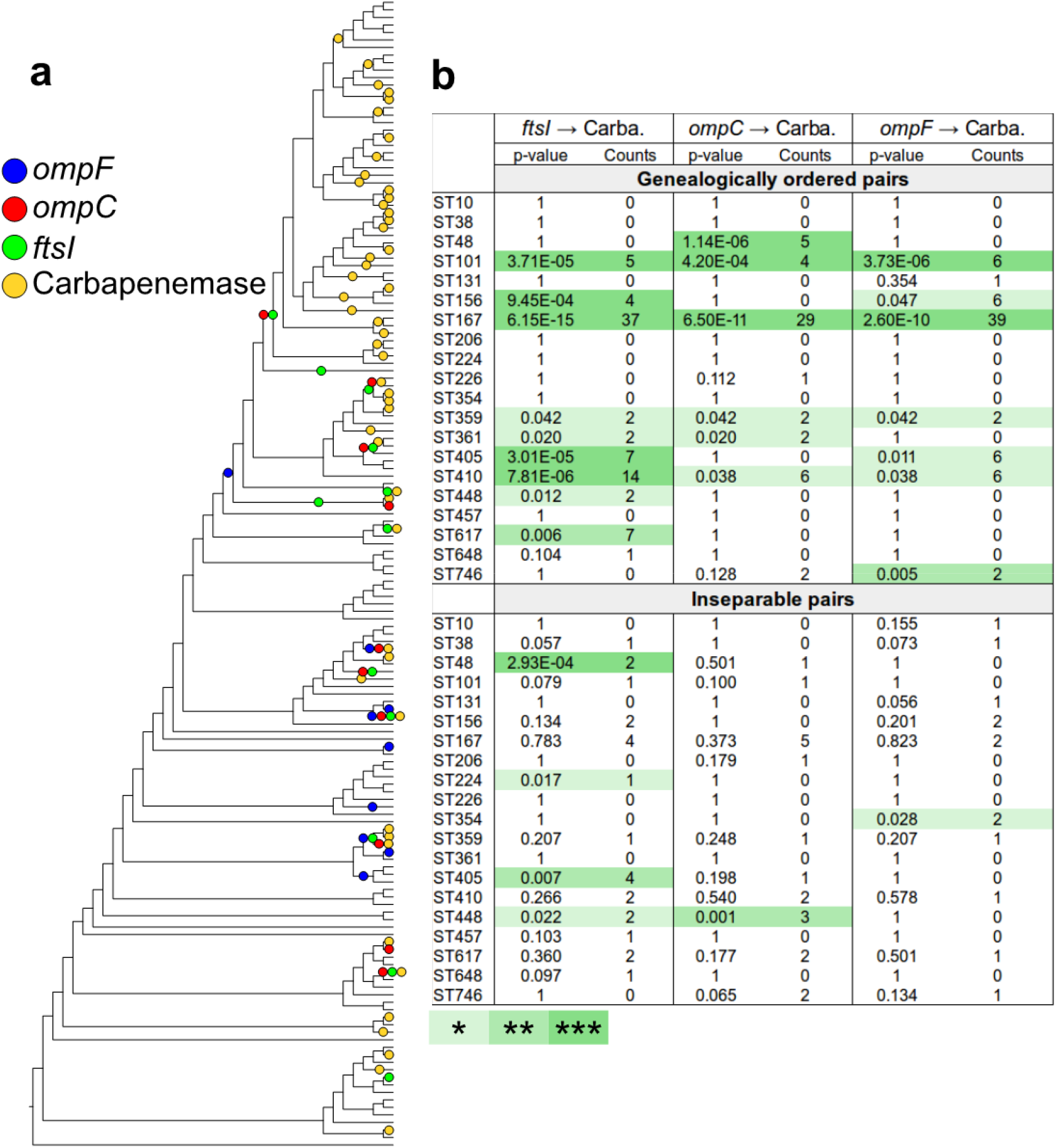
Test for the independence of the acquisition of carbapenemase alleles in defined genetic backgrounds. **a**. Cladogram obtained from the maximum likelihood tree estimated for the ST167 *E. coli* isolates (Supplementary Fig. 3). Four different genetic events, represented by coloured circles, have been placed by parsimony on the tree. In blue, mutations affecting the *ompF* gene (inactivation, by frameshift of premature stop codons, and mutations in the regulatory region); in red, genetic events affecting the *ompC* gene (gains of new *ompC* alleles by homologous recombination, inactivation of the gene, and non-synonymous mutations modifying the charge of AA localized in the pore lumen; in green, homologous recombination of an *ftsI* alleles with a 4 codon insertion (Fig. 2); and in yellow, acquisition of a carbapenemase gene. **b**. Testing for the independence between the acquisition of carbapenemase genes and mutations in porin genes, and/or *ftsI*. *, P < 0.05; **, P < 0.01; ***, P < 0.001.

### Acquisition of carbapenemase genes was preferentially selected in backgrounds mutated in *ompC*, *ompF* and *ftsI*

The observation of frequent co-occurrences of mutations in the three genes and acquisition of a carbapenemase gene is indicative of a genetic dependency between these events. In order to statistically test for the independence of two events in the phylogeny of each ST, we applied the methods (EpiCs) developed by Behdenna et al.^28^. This method takes into account the topology of the tree and the node at which each event is predicted, by parsimony, to have occurred (Fig. 6a). The test is based on a probabilistic framework that computes the exact probability of counts of co-occurrences (two events in the same branch) and/or subsequent events (one preceding the other in the tree). This statistical analysis was repeated on the 20 STs containing at least four CP-*Ec* isolates after removing redundancy. In each case, both models, ie: mutations occurring first or carbapenemase gene being acquired first, were tested. We obtained no evidence for the model where carbepenemase genes were acquired first. In contrast, in 11 STs, a significant association was observed for mutations in *ftsI* and the acquisition of a carbapenemase gene, with the mutation predicted to have occurred first in nine ST. Similarly, *ompC* and *ompF* mutations show a significant association with carbapenemase acquisition in seven STs each. For both porin genes, mutations were predicted to have occurred first in six STs. In three STs, mutations in the three genes precede the acquisition of the carbapenemase gene: ST167, ST101 and ST359. This result showed that carbapenemase genes within these 11 ST are preferentially acquired in a genetic background with a reduced susceptibility to β-lactams resulting of mutations in these three genes.

**Fig. 6.**
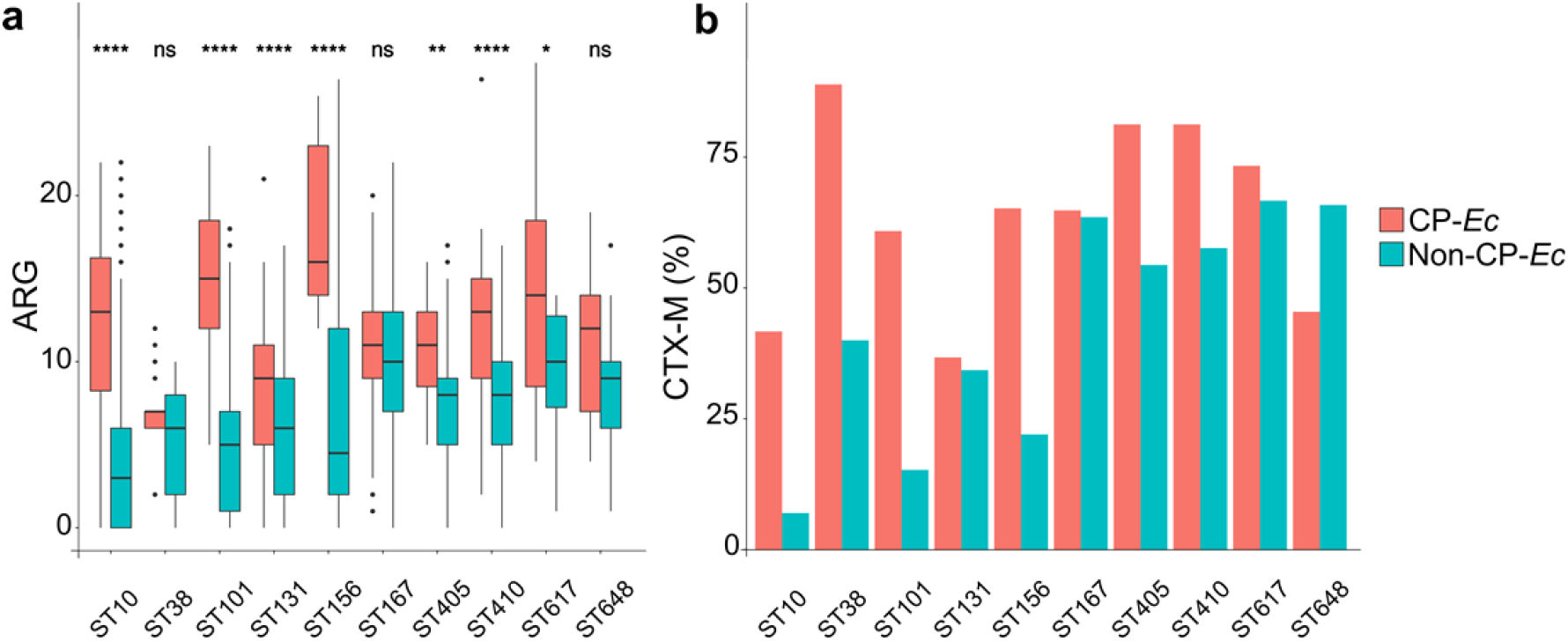
Occurrence of ARG and *bla*_CTX-M_ genes in CP-*Ec* isolates. a. Comparison of the number of ARG between CP-*Ec* and non-CP-*Ec* for the 10 STs encompassing more than 10 CP-Ec isolates. The horizontal lines in the boxes represent the median number of ARG. The box boundaries represent the first and third quartiles of the distribution and box-plot whiskers span 1.5 times the interquartile range of the distribution. Outliers are denoted as black points outside whiskers. Statistical significances were tested with a one-sided Wilcoxon rank-sum test. *, P < 0.05; **, P < 0.01; ****, P < 0.0001; ns, non-significative. b Comparisons in % of the presence of *bla*_CTX-M_ genes between CP-*Ec* and non-CP-*Ec.*

We did not detect such an association for ST131 (49 nr-CP-*Ec* isolates), ST10 (12 nr-CP-*Ec* isolates), ST648 (11 nr-CP-*Ec* isolates), ST226 (7 nr-CP-*Ec* isolates) and ST38 (26 nr-CP-*Ec* isolates). ST648 and ST38 belong to phylogroup D, the phylogroup that served as the source for the dissemination of specific *ompC* alleles by recombination in other CP-*Ec* lineages. As these alleles were present in the ancestors of the two STs and did not result from mutation or recombination, they were not considered in our association analysis although they probably confer a predisposition to acquire a carbapenemase gene. ST10 was the most numerous ST analysed in this study with 528 nr-isolates and showed a low rate of CP-*Ec* of 2% (Supplementary Fig 8). Despite the large number of CP-*Ec* ST131 isolates, none showed a four codons insertion in *ftsI* or an AA change in OmpC predicted to decrease susceptibility. In addition, among the 29 ST131 isolates with an inactivated *ompF* gene, only eight carry a carbapenemase gene. Furthermore CP-*Ec* were equally distributed in the four ST131 lineages A, B, C1 and C2 (Supplementary Fig. 7). Therefore, acquisition of carbapenemase genes in ST131 isolates might proceed according to a different path.

### Carbapenemase genes were more frequently acquired in MDR backgrounds

A characteristic of the ST410 OXA-181 subclade compared to other ST410 *Ec* isolates is a globally higher number of ARGs. To determine whether this observation can be extended to other CP-Ec isolates, we systematically analysed for their ARG content the isolates belonging to the 10 STs with more than 10 CP-Ec isolates. In most STs CP-*Ec* isolates showed a significantly higher number of ARG than non-CP *Ec* isolates. Only in ST38, ST167, and ST 648 the number of ARG was not significantly superior in CP-*Ec* (Fig. 6). Note that both CP-*Ec* and non-CP-*Ec* ST167 isolates show a high number of ARG (median=10). Similarly, we observed a higher percentage of CTX-M enzymes among CP-*Ec* isolates compared to non-CP-*Ec* isolates except in ST131 and ST648 (from phylogroup D).

### Mutations in *ftsI*, *ompC* and *ompF* associated with CP-*Ec* contribute to decreased susceptibility to β-lactams

Our data show that specific mutations in *ompC*, *ompF* and *ftsI* have been frequently selected in lineages that thereafter acquired carbapenemase genes by LGT. To further decipher the consequences of these mutations, we tested experimentally their impact on the susceptibility of *E. coli* to β-lactams. We first determined the contribution of the three mutations in *ftsI* (YRIN insertion, E349K and I532L) identified in the OXA-181 lineage. To this aim we constructed derivatives of the ST10 *s*train MG1655 with combinations of these mutations (Fig. 2b). Individually, each mutation showed only a small effect on susceptibility to β-lactam targeting PBP3. However, the combination of two or three mutations led to a stronger decrease in the susceptibility to these antibiotics. In particular, the MG1655 PBP3 derivative with the three modifications showed, in the absence of any β-lactamase, a 32, 16 and 14-fold increase in the MIC to the third-generation cephalosporin cefotaxime, to the monobactam aztreonam and to piperacillin respectively. This strain showed a slight increase in the MIC to ertapenem (x4), which mainly targets PBP2 and to a lesser extend PBB3, but no difference in the MIC to meropenem and imipenem that show low affinity for PBP3^29^.

To test the impact of mutations and recombination in *ompC* on β-lactams permeability, we complemented an *E. coli* K12 strain lacking the two major porins and carrying pOXA-232 (a high copy number plasmid encoding the carbapenemase *bla*_OXA-232_^30^) and tested for susceptibility to β-lactams (Fig. 3c). The wild type (wt) ST167 (CC10) *ompC* allele, and its G137D derivative, the ST38 (phylogroup D) allele, the wt ST410 allele and its R195L derivative were cloned into the medium copy number vector pACYC184^24^. Complementation with the different alleles of *ompC* led to an increased susceptibility to most β-lactams tested. However, we observed a differential effect of the different alleles of *ompC* (Fig. 3c). In particular we observed that strains expressing the R195L, the G137D and the ST38 *ompC* alleles showed reduced susceptibilities to cefalotin, cefoxitin, moxalactam and ertapenem compared to the strain complemented with the wt ST167 and ST410 alleles. These results confirm our prediction that the two *ompC* variants and the ST38 allele associated with Cp-*Ec* isolates show lower permeability towards different ß-lactams including ertapenem than their wild type counterparts.

In *E. coli*, the OmpF porin has been shown to contribute to the permeation of β-lactams into the periplasm and to susceptibility to these antibiotics^31^. To estimate the impact on β-lactam susceptibility of the mutation in the *ompF* promoter region identified in the ST410 OXA-181 subclade we quantified *ompF* mRNA by qRT-PCR. We compared *ompF* at normal and high osmolarity (LB and LB supplemented with 1M NaCl) between two isolates from the OXA-181 subclade (mutated) and two isolates from the FQR clade (non-mutated). As control, we also quantified *ompC* expression. We observed a 15 to 30 and 5-fold reduction in *ompF* expression in LB and LB-NaCl respectively in the mutated isolates compared to wild-type, whereas *ompC* expression remained unchanged (Fig. 2d). This confirmed that the regulatory mutation identified in the OXA-181 subclade leads to a decreased *ompF* expression in these isolates that will reduce entry of β-lactams into the periplasm and antibiotic susceptibility.

### Isolates of the OXA-181 subclade show higher resistance without *in vitro* fitness cost as compared to other OXA-181 *Ec* isolates of the FQR clade

Independently, mutations in *ftsI* and *ompC* affected the susceptibility to β-lactams of a laboratory strain. To determine the impact of mutations and ARG on β-lactam resistance and fitness in ST410 isolates, we analysed the phenotype of five *bla*_OXA-181_ *bla*_CTX-M-15_ *Ec* clinical isolates from different lineages of the ST410 FQR clade (Supplementary table 8) in comparison with an isolate expressing only CTX-M-15: two isolates from the OXA-181 subclade mutated in *ompC*, *ompF* and *ftsI* (YRIN insertion), one isolate mutated in *ftsI* (YRIK insertion) and the two other isolates without any mutation in the three genes. We observed, by disk diffusion assay and Etest, a gradual decrease in the susceptibility to diverse β-lactams between the four groups of isolates: CTX-M15 < CTX-M15, OXA-181 < CTX-M15, OXA-181, YRIK insertion in PBP3 < OXA-181 subclade (Fig. 7a and b). According to CLSI breakpoints, isolates from the OXA-181 subclade were resistant to almost all β-lactams tested except doripenem and imipenem and intermediate to meropenem and mecillinam. Higher resistance was partly due to differences in β-lactamase gene content including *bla*_OXA-181_ (Supplementary table 8). However, mutations in *ftsI*, including the YRIK and YRIN insertions were likely responsible for the decreased susceptibility of ST410 *Ec* isolates to β-lactams targeting PBP3 like ceftazidime and aztreonam (Fig. 7b). The isolates from the OXA-181 subclade showed a decreased susceptibility to ertapenem and meropenem likely resulting from *ompC* and *ftsI* mutations for ertapenem.

**Fig. 7.**
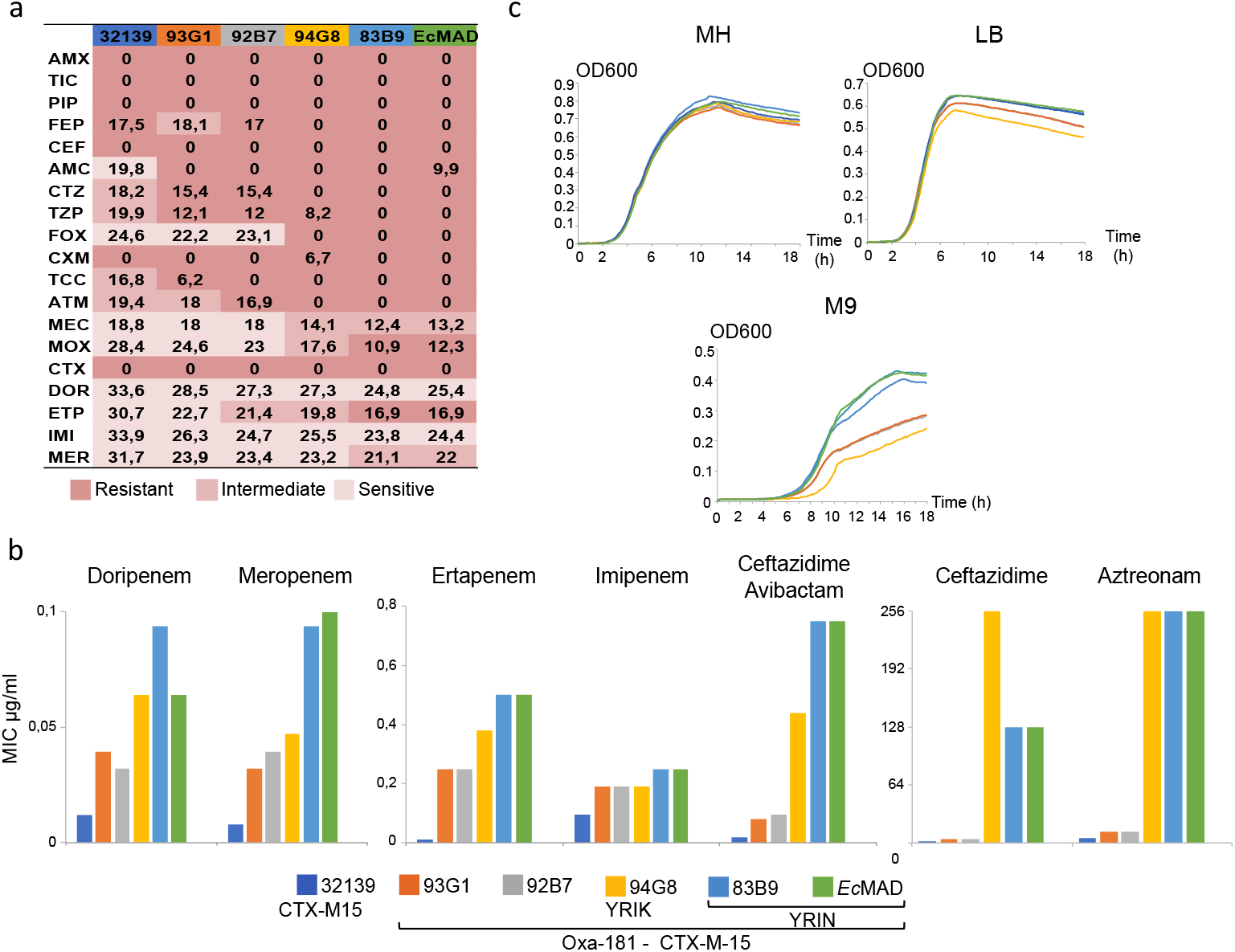
β-lactam susceptibility profiles and fitness of *Ec* ST410 strains. **a.** β-lactam susceptibility determined by disk diffusion. Diameters are indicated in mm. Resistant, Intermediate and Sensitive according to CLSI guidelines^32^ are indicted by colours as defined in the figure keys. 32139 carries *bla*_CTX-M15_, 93G1 and 92B7 carry *bla*_CTX-M15_ and *bla*_OXA-181_, 94G8 carries *bla*_CTX-M15_ and *bla*_OXA-181_ and a mutated *ftsI* gene (YRIK), 8389 and *Ec*Mad belong to the OXA-181 subclade. ****b.**** Minimal inhibitory concentrations (MIC) determined by E-test for selected β-lactams. **c.** Growth curves in rich (LB and Müller Hinton, MH) and minimal (M9) medium. Curves represent the average value of 10 experiments. Box plot representations of the area under the curve of replicates determined by using growthcurver ^33^ are given in Supplementary Fig. 9. Abbreviations: AMX, Amoxicillin; TIC, Ticarcillin; PIP, Piperacillin; FEP, cefepime; CEF, cephalothin; AMC, Amoxicillin clavulanic acid.; CTZ, Ceftazidime; TZP, piperacillin-tazobactam; FOX, Cefoxitine; CXM, Cefuroxime; TCC, Ticarcilline-clavulanic acid; ATM, Aztreonam; MEC, Mecillinam; MOX, Moxalactam; CTX, Cefotaxime; DOR, Doripenem; ETP, Ertapenem; IMI, Imipenem; MER, Meropenem. Antibiotic resistance gene repertoires of the six strains are given in Supplementary table 8.

To determine whether these mutations have an impact on fitness we compared growth parameters in LB, MH and M9 media as proxy. Despite the higher resistance to antibiotics, we did not detect any significant differences in the growth parameters in rich medium among the six isolates tested (Fig. 7c) suggesting these mutations have no fitness cost or their effect has been compensated by other mutations. The two isolates from the OXA-181 subclade and the non-OXA-181 isolate 32139 grew at a higher OD600 in minimal medium than the three other isolates (Fig. 7c). Therefore, the high level of resistance to most β-lactams and the decreased susceptibility to carbapenems of isolates of the OXA-181 subclade do not seem to be at the expense of a lower *in vitro* fitness for the isolates we have analysed.

## Discussion

During the past twenty years the increased prevalence of ESBL-producing enterobacteriaceae led to an increased use of carbapenems and selection of CP-*Ec* strains. The dissemination of Cp-*Ec* lineages is particularly feared. The global dissemination of antibiotic resistant clones and their evolutionary trajectory result from a trade-off between acquired resistance and biological cost in the absence of antibiotic^34^. However, how this trade-off is reached during *in vivo* evolution is largely unknown as it likely depends on multiple factors, such as antibiotic usage, which differs between areas of the world^35^. Also, the same clone may spread across various human and non-human sectors and successively undergo highly different selective pressures. In that context both the surveillance of emerging Cp-*Ec* clones in a One Health perspective and their thorough genetic analysis are required to characterize their evolutionary trajectories and prevent dissemination of other and possibly more virulent clones.

Here, we first analysed the worldwide disseminated CP-*Ec* OXA-181 ST410 subclade and extended this analysis to the whole *E. coli* species. We showed that carbapenemase genes were preferentially acquired in lineages already mutated in three genes contributing to β-lactam resistance: *ompC*, *ompF* and *ftsI*. Indeed, in 13 STs representing 54% (n=234) of the non-redundant CP isolates analysed in this work, a combined phylogenetic and statistical analysis revealed a significant association between mutations of these genes and subsequent acquisition of a carbapenemase gene (Fig. 5). Globally we found that 45%, 32% and 38% of the studied CP-*Ec* isolates were mutated in *ftsI*, *ompC* and *ompF* respectively. The association was best exemplified by a large clade among ST167 isolates first defined by a mutation in an OmpR box within the *ompF* promoter region. Within this clade, six events of recombination led to the acquisition of mutations in *ftsI* and four genetic events modified the *ompC* gene: two recombination events leading to its replacement by an allele from a phylogroup D strain and two homoplasic G137D mutations affecting AA of the pore lumen. Eventually, multiple events of acquisition of carbapenemase genes were selected (Supplementary Fig. 3). Interestingly, the encoded carbapenemases belonged to different families that differ by the levels of carbapenem resistance and by the spectrum of β-lactams they hydrolyze^36^. Therefore, the scenario originally detected in the ST410 OXA-181 subclade expands to numerous other STs and does not seem to depend on the carbapenemase family.

Such a situation is reminiscent of what has recently been observed in the *K. pneumoniae* high-risk clones ST258, ST512 and ST11^37^. In these clones, the acquisition of carbapenemase genes was frequently associated with the inactivation of the porin gene *ompK35* (equivalent of *E. coli ompF*) and mutations in *ompK36* (equivalent of *ompC*). In CP-*Ec*, we observed *ompC* inactivation in a few isolates, probably due to a high fitness cost of this event. Instead, like in *K. pneumoniae*, the selected mutations affected *ompC* permeability towards β-lactams (Fig. 3c) while probably keeping the global function of the porin. This likely leads to a lower fitness cost than gene inactivation, as also hypothesized in *K. pneumoniae* ^37^. In contrast, both *ompF* inactivation or mutations with likely a lower fitness cost such as those in its promoter region were observed in CP-*Ec*. This indicates that, under *in vivo* growth conditions, the OmpF porin might be more easily dispensable than OmpC. On the other hand, PBP3 is essential for cell division and selection of *ftsI* mutants is expected to be highly evolutionary constrained. This might explain the extremely rare *ftsI* mutations previously reported in clinical lineages of *E. coli*. This contrasts with the high frequency of *ftsI* mutations observed among Cp-*Ec* isolates.

During the evolution of CP-Ec lineages, recombination events involving *ftsI* and *ompC* were pervasive throughout the *E. coli* species except among phylogroup B2. We identified four combinations of mutations in *ftsI* frequently associated with CP-*Ec* isolates (Fig. 2a). To our knowledge these modifications are the only mutations contributing to β-lactams reported in natural *E. coli* isolates^19,25^. Nevertheless, the A413V mutation found in association with three types of insertions, has recently been selected during serial passages of *E. coli* in the presence of aztreonam and shown to lead to a four-fold decreased susceptibility to this antibiotic ^38^. This result is in agreement with our prediction that this mutation might contribute to resistance to β-lactams of isolates carrying YRIK, YTIP or YRIP insertions as we have shown for the two SNPs associated with the YRIN duplication (Fig. 2b). The chance of mutation combinations leading to a significant decrease in susceptibility is likely very low, but the selective advantage strong. In agreement with this hypothesis, the phylogenetic reconstruction of the recombination events showed that these combinations arose likely only once and disseminated widely across the *E. coli* species by LGT, as we identified 48 events of recombination (Fig. 4). Most of these recombination events were associated with at least one CP-*Ec* isolates (n=46) and in 24 cases it corresponded to disseminated lineages (i.e. with more than three isolates from different geographical origins). We observed a similar situation for the porin OmpC, with 20 events of recombination events of *ompC* alleles originating from phylogroup D isolates. The chromosomal region next to the *ompC* gene has been shown as a hotspot of recombination ^39^. However, we observed that the acquisition of this specific allele was associated with acquisition of carbapenemase genes in most cases (n=15). Recombination has been shown to play a major role in β-lactam resistance in pneumococcus^40^ or in *Neisseria spp*^41^. Altogether, our data show for the first time that in addition to the LGT of β-lactamase genes located on MGE, recombination has a significant contribution in β-lactam resistance in *E. coli*, including carbapenems.

Further comparisons of β-lactam susceptibility and fitness among ST410 isolates carrying the same *bla*_OXA-181_-bearing plasmid and *bla*_CTX-M-15_ gene with different patterns of mutations in *ompC*, *ompF* and *ftsI* showed that in these clinical isolates and particularly in the broadly disseminated OXA-181 subclade, these mutations were not associated with a fitness cost (Fig. 7c). We also observed that increased resistance to β-lactams could be attributable to mutations in the three genes, in agreement with our experimental study of these mutations individually (Fig. 2 and 3). In particular, for the OXA-181 subclade, we observed an additional decrease in the susceptibility to ertapenem. Interestingly, ertapenem shows a higher biliary excretion than other carbapenems and was found to have a stronger impact on the intestinal microflora^42^. *ftsI* mutations were also found to be selected during *in vitro* evolution in the presence of ertapenem but not meropenem^43^. However, other β-lactams like aztreonam might also have contributed to the selection of these combinations of mutations^38^.

A characteristic of the ST410 OXA-181 subclade is a higher number of ARG than in other ST410 *Ec* isolates (Supplementary Fig. 1), suggesting that antibiotic pressure was a major contributor to the evolution of this clone. In particular, the phylogenetic analysis reveals that mutations in *ompC*, *ompF* and *ftsI* occurred in a context already resistant to fluoroquinolone and expressing CTX-M-15 ESBL (Fig. 1). A systematic analysis of the number of ARG across the *E. coli* species showed that a higher number of ARG in CP-*Ec* compared to non-CP-Ec isolates was a common feature in all STs with more than 10 CP-*Ec* isolates. A single exception was ST167 where a similarly high number of ARG was observed in both CP and non-CP-*Ec* isolates analysed (Fig. 6). Therefore, for these STs, acquisition of carbapenemase genes occurred more frequently in an MDR background. We also observed a frequent co-occurrence of CTX-M family ESBL and carbapenemase genes in most ST, but not in the ST131 lineage. In all, these data suggest a long-term and step by step evolution of those lineages with episodic periods of selection and dissemination. As a first step, specific mutations in *ompF, ompC* or *ftsI* would have been fixed in isolates already expressing different β-lactamases including ESBL enzymes, leading to low levels of resistance to carbapenems^44^. This scenario is compatible with our phylogenetic analysis and the high frequency of *bla*_CTX-M_ genes in isolates mutated in *ftsI* (78% of nr-isolates) and with our experimental data on the effect of mutations in porin genes and in *ftsI* on β-lactam susceptibility (Fig. 2 and 3). In a second step, under antibiotic pressure, the combination of these mutations and β-lactamase expression might have favored the efficient conjugative transfer of plasmids carrying carbapenemase genes from other CP-bacterial species by increasing the proportion of donor and receptor bacteria^45^. This might have occurred in the context of low levels of carbapenems or other β-lactams, such as found in the gut during parenteral administration of antibiotics with biliary excretion. This model could also explain the high prevalence of ST38 isolates observed both in England and in France^8,9^, as 24 out of 27 (89%) ST38 CP-*Ec* isolates express a CTX-M class enzymes. The specific ST38 *ompC* allele with reduced permeability to different β-lactams including ertapenem together with CTX-M ESBL would have favored the fixation of carbapenemase genes.

Although CP-*Ec* were frequent among the ST131 isolates studied here, with 49 nr-isolates (66 in total), we did not observe any case of four AA insertions in *ftsI* or of mutations affecting AA in the pore lumen of OmpC among the 402 nr-genome sequences we have analysed. Furthermore, CP-*Ec* isolates were broadly distributed among the different ST131 lineages and were not associated with CTX-M type ESBL since only 37% expressed also β-lactamase of this class (Fig. 6b). The selection for ST131 CP-*Ec* isolates might therefore follow a different path compared to others CP-*Ec*, which might be related to their higher and human-specific pathogenicity^46^. ST131 CP-*Ec* isolates might arise sporadically in patients following conjugation of carbapenemase gene carrying plasmids from another CPE and subsequent selection by β-lactams, including treatments with carbapenems. These transconjugants would be more frequently detected in clinics due to their high pathogenicity. On the other hand, possibly due to their lower fitness they would not disseminate globally. Indeed, despite the high prevalence of ST131 CP-*Ec* reported in different studies^11,47,48^, there was no indication of global dissemination of a specific lineage among the ST131 CP-*Ec* genome sequences we have analysed (Supplementary Fig. 7).

## Material and methods

### Bacterial isolates, growth conditions and antibiotic susceptibility testing

Features of the clinical *E. coli* isolates analysed in this work are listed in Supplementary table 1. 50 *Ec* ST410 isolates came from the strain collection of the Fr-NRC for Antibiotic Resistance. Four *Ec* ST410 clinical isolates came from the Microbiological collection of the Public Health Faculty of the Lebanese University (Tripoli, Lebanon) and three ST410 isolates of animal origin from the ANSES strain collection. Test for OmpC permeability to β-lactam was performed in strain W3110 deleted for *ompC* and *ompF* genes^49^. Antibiotic susceptibility was performed by the disk diffusion method following the Clinical & Laboratory Standards Institute (CLSI) guidelines^32^ or by Etest (Biomérieux) following the manufacturer’s recommendations. For W3110 Δ*ompC* Δ*ompF* pOXA-232 strains carrying pACYC184 derivatives, disk diffusion assays were performed on Mueller Hinton agar plates supplemented with 2mg/l chloramphenicol. Fitness was determined by growth curve analysis with an automatic spectrophotometer Tecan Infinite M200 during 24 hours in LB, Mueller Hinton or M9 media supplemented with 0.4% glucose. Growth metrics were estimated with the R package “growthcurver” ^33^. The area under the curve (AUC) which include contributions of the most important growth parameters (log phase, growth rate, and the carrying capacity) was used as a growth metric.

### Genome sequencing, and genome sequences retrieved from sequence libraries

*Ec* genomes were sequenced by using the Illumina HiSeq2500 platform, with 100 nucleotides (nt) single reads for the four isolates from Lebanon and 100 nt paired-end reads for the other isolates. Libraries were constructed by using the Nextera XT kit (Illumina) following the manufacturer’s instructions. The OXA-181-producing *Ec*-MAD ST410 isolate was selected as a reference strain and sequenced to completion by using the long read PacBio technology. 10 947 *E. coli* and 1 451 *Shigella* genome sequences deposited in the NCBI database (19th June, 2018) were retrieved for a global analysis of the specificity of CP-*Ec* (Supplementary table 2). 96 additional *Ec* ST167 isolates were retrieved from Enterobase (https://enterobase.warwick.ac.uk/). Raw reads from 62 *Ec* ST410 and 21 *Ec* ST38 isolates identified in Enterobase (march 2017) were retrieved from the NCBI database. (Supplementary table 2). Redundancy in the genome collection was removed by filtering for isolates from the same study, diverging by less than 7 SNPs. We kept one randomly selected isolate. In case of differences in resistome, assuming that ARG loss was more likely than ARG gain, we kept an isolate with the largest number of ARG.

### Sequence assembly, genome annotation and mutation identification

The PacBio reads were assembled with the RS_HGAP_Assembly.3 protocol from the SMRT analysis toolkit v2.3^50^, and with Canu^51^. The consensus sequence was polished with Quiver^50^ and by mapping Illumina reads. Illumina sequenced isolates were assembled with SPAdes^52^, and the quality of the assemblies was assessed with Quast^53^. Contigs shorter than 500 bp were filtered out. All assemblies were annotated with Prokka^54^. The presence of antibiotic resistance genes, and plasmid replicons was assessed with ResFinder^55^ and PlasmidFinder ^56^ respectively. The software was run in local from the script and databases downloaded from the repositories of the Centre for Genomic Epidemiology (https://bitbucket.org/genomicepidemiology/). Graphs of genomic regions of interest were drawn with genoplotR^57^. All downloaded genomes were reannotated with prokka^54^. For each ST type analysed (Warwick scheme) the pangenome was characterized with Roary^58^, and the AA sequences of OmpC, OmpF, GyrA, ParC, and FtsI were identified from the orthologous table generated by Roary. To determine the presence of different *ompC* alleles within the different STs, OmpC AA sequences were clustered with cd-hit^59^, with a sequence identity threshold of 0.95. AA sequences for GyrA and ParC were aligned with the mafft L-INS-i approach^60^, and the amino acids at the QRDR positions for both genes were filtered with customized Perl scripts.

### Phylogenetic reconstruction

A core genome alignment was obtained for all *Ec* ST410 isolates, and strain 789 that belongs to the ST88 as an outgroup with parsnp^61^. Non-recombinant SNPs were extracted from the core genome alignment with Gubbins^17^, and the Maximum likelihood (ML) phylogeny was obtained with RAxML^16^, using the General Time-Reversible (GTR) substitution model with a gamma-distributed rate. The same procedure was followed for the phylogenetic reconstruction of the different *E. coli* lineages. Maximum likelihood phylogeny of OmpC protein sequences was inferred with RAxML^16^. OmpC protein sequences were aligned with the mafft L-INS-i approach^60^. Gblocks^62^ was used to “clean” the alignment and the best fit model (WAG, with a gamma distribution) was estimated with protest 3^63^. The visual display of phylogenetic trees was done with FigTree (http://tree.bio.ed.ac.uk/software/figtree/), and annotated trees with the script plotTree.R (https://github.com/katholt/plotTree). The final version of the tree was edited with Inkscape (https://inkscape.org/es/).

### Mapping, variant calling, and identification of SNPs of interest

Sequence reads for *Ec* ST410 isolates were mapped against the complete reference genome *Ec*-MAD with BWA^64^. Variant calling was performed with the Genome Analysis Toolkit v 3.6.0^65^. The criteria for variants were: occurrence of the alternative base in more than 90% of the reads covering the position, a depth coverage of at least 10 (DP > 10), a quality by depth (QD) > 2, a fisher strand bias (FS) < 60, a mapping quality (MQ) > 40, a mapping quality rank sum test (MQRankSum) > −12.5, and a read position rank sum test (ReadPosRankSum) > −8. Variants associated with different clades of the *Ec* ST410 phylogeny were extracted with vcftools^66^, and finally they were annotated with snpEff^67^. The effect of the non-synonymous mutations was assessed with the Sorting Intolerant From Tolerant (SIFT) algorithm^20^. The algorithm searches for protein homologs in the refseq database using mutated proteins as query and assigns a score to each position. This score is weighted by the properties of the AA changed. If this score is below a threshold (0.05) the change is predicted to be functional.

### Testing the independence between mutations in *ftsI*, *ompF*, and *ompC*, and the acquisition of carbapenemase genes

To assess the association between the different genetic events we used the method developed by Behdenna et al.^28^ implemented in the software EpiCs. The described events were mapped on the tree by parsimony, and the probability distribution of the number of paired events occurring in the tree was computed under the null model of independence. Two types of paired events are described in the method: inseparable pairs, when both events occur in the same branch, and genealogically ordered pairs, when the second event is found in a node more recent than the first one. We have considered the following genetic events: (i) “*ompC* mutations” encompassing acquisition of *ompC* alleles from phylogroup D strains by recombination, *ompC* mutations changing the charge of AA in the pore lumen, and *ompC* inactivation; (ii) “*ompF* mutations” including *ompF* gene inactivation and point mutations in the OmpR binding sites of its promoter; (iii) “*ftsI* mutations” including the four different four-codon duplications (YRIN, YRIK, TYPI, and YTIP) in *ftsI* and (iv) the acquisition of a carbapenemase gene. We have focused our analysis on the independence between carbapenemase gene acquisition and mutation in each of the three genes, ompC, *ompF* and *ftsI*.

### Complementation of the W3110 Δ*ompC* Δ*ompF* strain

*ompC* alleles and their regulatory regions were cloned into the medium copy number vector pACYC184^24^ following amplification by primers *ompC*_*Xba*_F and *ompC*_*Hind*_R (Supplementary Table 9), digestion by *Xba*I and *Hind*III restriction enzymes and ligation into the vector digested by the same enzymes. Ligation was transformed into commercial *E. coli* TOP10 competent cells (Invitrogen). The absence of mutation was checked by Sanger sequencing. Plasmid containing the *ompC* gene from MG1655 or the G137D mutated allele, the ST410 wt allele and the R195L mutated allele from *Ec*MAD and the ST38 allele as well as the empty vector were introduced into competent W3110 Δ*ompC* Δ*ompF* pOXA-232. Competent cells were prepared by the CaCl_2_ method^68^. Plasmid pOXA-232^30^ was prepared from an ST231 *Klebsiella pneumoniae* isolate from the Bicêtre hospital collection carrying this plasmid. Plasmid content in transformants was verified by plasmid DNA extraction (Qiagen) and Sanger sequencing.

### Construction of *ftsI* mutant strains

The three mutations identified in the *ftsI* gene of the MAD strain were reconstructed in a MG1655 genetic background to analyse their effects on antibiotic resistance. To this aim we introduced the 12 nt insertion (YRIN form) and the two non-synonymous SNPs (E349K and I532L) into the *Ec* strain MGF (MG1655strepR_F’tet_ΔtraD::Apra) by TM-MAGE^69^. Briefly, overnight culture of strain MGF transformed by pMA7SacB was used to inoculate 5 ml LB medium supplemented with tetracycline (7.5 μg/l) and carbenicillin (100 μg/l) (LB-TC) and grown at 37°C until OD_600_ reached 0.6-0.7. The recombinase and Dam methylase were induced by addition of L-arabinose (final concentration of 0.2% w/v), and further incubation for 10 min. Cultures were then chilled for 15 min on ice and centrifuged at 7300g at 4°C. Two successive washes by 50 and 10 ml of cold water were performed and the final pellet was resuspended in 200 μl water. 100 μl of cells were used for electroporation with 2 μl oligonucleotides Mut1ftsI or Mut2ftsI (Supplementary table 9) alone or in combination at 20 μM each. The Mut1f*tsI* oligonucleotide carries both the 12 nt insertion and the E349K mutation, while the Mut2*ftsI* oligonucleotide has the I532L mutation. The content of the electroporation cuvette was used to inoculate 5 ml of LB-TC and submitted to three additional cycles of growth-induction-preparation of electrocompetent cells and electroporation. Following the last electroporation step, cells were resuspended in1 ml LB and plated onto LB-TC agar plates. Mutations in isolated colonies were tested by PCR using primers complementary to mutant or wild-type alleles (Supplementary table 9). Mutated colonies were grown on plates containing 10 g/l tryptone, 5 g/l yeast extract, 15 g/l agar and 5% w/v sucrose for plasmid curing. Mutant strains were sequenced by using Illumina MiSeq platform, with 150 nt paired-end reads and Nextera XT kit (Illumina) for library preparation. Reads were mapped onto the MG1655 genome (Genbank NC_000913.3) to confirm that mutations in *ftsI* gene have been correctly introduced and to check that the rare off target mutations are not predicted to interfere with the β-lactam susceptibility phenotype (Supplementary table 10).

### RNA extraction and quantitative RT-PCR

Bacteria were grown in LB medium until OD600 reached 0.30-0.33, an osmotic shock was applied by supplementing 10 ml of culture with 0.3M, final concentration of Sodium Chloride (NaCl) or with the same volume of water as control and further incubation for 20 minutes. Bacterial pellets were collected and stored at −80^⍛^C. Total RNA was extracted with the Total RNA purification kit Norgen Biotek. cDNAs were synthetized from 500 ng of RNA with the Superscript II reverse transcriptase (Invitrogen, Life Technologies). Primer pairs were designed for the *ompC* and *ompF* genes, targeting divergent regions from these two genes, and for the reference gene *recA* (Supplementary Table 9). The SYBR Green PCR kit (Applied Biosystems, Life Technologies) was used to perform quantitative PCR, and the relative expression of porin genes was measured by a standard curve method where the regression analysis was performed from serial dilutions of a mixture of control cDNAs. The expression value of each gene was normalized against the expression of the housekeeping gene *recA*. Each point was measured in triplicate, and three independent cultures were used for each strain in each condition.

### Statistical analysis

The statistical significance of the differences in expression in RT-PCR experiments was assessed by using a two-tailed t-test. The statistical significance of the differences in the number of ARG between bacterial groups in different STs was assessed using the Wilcoxon rank sum-test implemented in R (v3.4.4). One-sided test was used for the comparison of the number of ARG, and two-sided test was used for the comparison on the area under the curve between six isolates from the ST410 FQR clade.

### Data availability

Illumina reads from the 57 newly sequenced isolates and the complete genome assembly of strain Ec-MAD have been deposited in the EMBL nucleotide sequence database (http://www.ebi.ac.uk/ena) under study accession number PRJEB27293 and PRJEB27274 respectively. The accession numbers for individual isolates are listed in Supplementary Table 1. Code of software scripts could be obtained upon request.

## Acknowledgements

This work was supported by grants from the French National Research Agency (ANR LabEx IBEID and ANR-10-LABX-33), from the Joint Program Initiative on Antimicrobial Resistance (ANR-14-JAMR-0002) and from One-Health EJP ARDIG. The authors thank Alexandre Almeida, Eduardo Rocha and Adriana Chiarelli for their discussion and critical reading of the manuscript, Guillaume Achaz for advices for the EpiCs analyses, Sylvie Letoffe, and Laurence Ma for their help in performing the TM-MAGE and the Illumina sequencing respectively. They also thank Jean-Michel Pages for the gift of strain W3110 Δ*ompC*, Δ*ompF*.

## Authors contribution

RPN, IRC and NC performed experiments; LG, JT, JYM, MH, RB and TN collected and provided the samples for the study; RPN, retrieved genome sequences from public databases, wrote scripts and performed all the bioinformatics analyses; RPN, IRC, NC and PG analysed the data; TN provided expertise on CP-*Ec*; RPN and PG designed the study; PG was responsible for management of the project; PG and IRC built the evolutionary model; RPN, IRC, TN and PG wrote the manuscript. All authors approved the manuscript prior to submission.

## Conflict of interest

The authors declare no competing financial interests.

