## Supplemental Tables and Figures for "Stepwise evolution and convergent recombination underlie the global dissemination of carbapenemase-producing *Escherichia coli*"

.

Rafael PATIÑO-NAVARRETE, Isabelle ROSINSKI-CHUPIN, Nicolas CABANEL, Lauraine GAUTHIER, Julie TAKISSIAN, Jean-Yves MADEC, Monzer HAMZE, Remy A. BONNIN, Thierry NAAS[$] and [*]Philippe GLASER.

### Supplementary figure legends.

**Supplementary Fig. 1 Number of antibiotic resistance genes (ARG) according to the phylogenetic group among *E. coli* ST410 isolates.** The horizontal lines in the boxes represent the median number of ARG associated to the three clades: Basal (n=22); FQR clade (n=92) and OXA-181 subclade (n=41). The box boundaries represent the first and third quartiles of the distribution and box-plot whiskers span 1.5 times the interquartile range of the distribution. Outliers are denoted as black dots outside whiskers. Statistical significances were tested with a one-sided Wilcoxon rank-sum test. ***, P < 0.001.

**Supplementary Fig. 2. Recombination events in isolates of the *E. coli* ST410 FQR clade.** Phylogeny of the *E. coli* ST410 FQR clade (left). On the right side, the recombined blocks identified by Gubbins[1] are highlighted in red when they occurred in an internal branch and affect more than one isolate and in blue when they occurred in a terminal branch affecting a single isolate. The core genome of *E. coli* ST410 FQR strains is represented above the figure and annotated genes correspond to the limits of major recombination events.

**Supplementary Fig. 3. Phylogeny and mutations in non-redundant CP-*Ec* isolates of ST167, ST101 and ST156**. ML phylogenies were estimated as for Fig. 1, using 5,843, 21,810, and 18,746 non-recombinant SNPs for **a.** *Ec* ST167, rooted with the ST10 strain MG1655 (NC_00913) **b.** *Ec* ST101, and **c.** *Ec* ST156 respectively, both rooted with the ST1128 strain IAI1 (NC_011741). 14 distantly related non-CP-*Ec* ST156 isolates have been removed from the phylogenetic analysis. Branch tips indicates the presence and the type of carbapenemase according to the figure key on the left. On the right side of the tree are represented, from left to right: $bla_{\text{CTX-M}}$ ESBL, *gyrA* and *parC QRDR* mutations, mutations in the *ftsI* gene, and genetic events affecting *ompF* and *ompC* according to the figure key at the bottom. SNPs in the *dcw* region are represented by small vertical red bars. Genes from the *dcw* locus are indicated by arrows and the *ftsI* gene in red. Black arrowheads indicate isolates used as reference for SNPs mapping in the *dcw* gene cluster.

**Supplementary Fig. 4. Phylogeny and mutations in STs including at least two isolates mutated in *ftsI*.** Maximum likelihood phylogenies were estimated from 21,101 (ST48), 13,531 (ST206), 20,092 (ST224), 16,572 (ST359), 18,649 (ST361), 18,992 (ST448), and 1,849 (ST617) core non-recombinant SNPs. Isolates used to root the trees are at the bottom of each tree. Branch tips represent the carbapenemase (or absence) type. Columns on the right side of the trees represent from left to right: *bla*$_{CTX-M}$ ESBL type, mutations in *gyrA* and *parC* QRDR and in *ftsI* and genetic events affecting *ompC* and *ompF* according to the figure key on the right. When different alleles of *ompC* where acquired by recombination within the same ST, they are numbered.

**Supplementary Fig. 5 Phylogeny and mutations in non-redundant *E. coli* isolates of phylogroup D.** Maximum likelihood phylogeny of phylogroup D isolates of ST38, ST354, ST405, ST457 and ST648 were obtained from 39,701, 41,662, 40,630, 43,491, and 39,577 core non-recombinant SNPs respectively. Isolates used to root the trees are at the bottom of each tree. Branch tips represent the carbapenemase type encoded or the absence of a carbapenemase according to the figure key. The columns represent, from left to right: *bla*$_{CTX-M}$ ESBL type, mutations in *gyrA* and *parC* QRDR, *ompC* and *ompF* according to the figure key.

**Supplementary Fig. 6. Recombination events at the *dcw* and *ompC* loci. a.** Recombination of the *dcw* cluster encompassing *ftsI* carrying an YRIN insertion. The *ftsI* gene (in green) of an *E. coli* ST101 isolate was affected by a 12-nucleotide duplication leading to an YRIN insertion and by two additional mutations (E349K and I532L). A 29.5 kb chromosomal region of an ST101 isolate mutated in *ftsI* comprising most of the *dcw* gene cluster was transferred by recombination into a CC10 strain, which transferred a 124-kb fragment containing the *dcw* locus to the common ancestor of the OXA-181 ST410 subclade. **b.** Recombination of *ftsI* carrying an YRIP insertion. A 65-kb region from an *Ec* ST156 isolate affected by YRIP duplication and A413V mutation was transferred by recombination into an *E. coli* ST224 strain. **c**. Recombination of *ompC* gene from an ST38 strain. A 22.7-kb region encompassing the *ompC* gene (in orange) was transferred by recombination into the MRCA genome from the ST167 clade of isolates expressing different carbapenemase genes (Supplementary fig. 3). Vertical black arrows indicate the limits of the recombined regions and green arrows the direction of exchange. Small vertical red bars represent SNPs compared to the donor strains. CDSs are indicated by blocks, upper line transcribed in the rightward orientation and lower line in the opposite orientation. The *dcw* gene cluster is in grey.

**Supplementary Fig 7. Phylogeny and mutations in non-redundant *E. coli* ST131 CP-*Ec* isolates.** Maximum likelihood phylogeny was obtained from the alignment of 48,705 non-recombinant SNPs. Tree was rooted with the ST405 strain Z1002 as an outgroup. Vertical lines represent ST131 clades (A, B, C1, C2) as previously described[2]. Branch tips represent the carbapenemase (or absence) type as defined in the figure key on the left. Columns represent, from left to right: *bla*$_{CTX-M}$ ESBL type, mutations in the QRDR regions of *gyrA* and *parC*, and mutations inactivating *ompC* and *ompF* porin genes as defined in the figure key on the right.

**Supplementary Fig. 7 Phylogeny and mutations of ST10, ST226, and ST746 non-redundant *E. coli* isolates.** Maximum likelihood phylogenies of ST10, ST226, and ST746 were estimated from 20,645, 10,041, and 69,372 core non-recombinant SNPs respectively. Isolates used to root the trees are at the bottom of each tree. In the ST10 final alignment the number of sequences was reduced from 765 to 137 by clustering sequences of non CP-*Ec* isolates with an identity over 99 % using cd-hit[3]. Branch tips represent the carbapenemase (or absence) type as defined in the figure key. Columns represent, from left to right: $bla_{\text{CTX-M}}$ ESBL type, mutations in *gyrA* and *parC* QRDR, *ompC* and *ompF* according to the figure key on the right.

**Supplementary Fig. 9. Estimated fitness of *E. coli* ST410 strains in rich and minimal media.** Fitness was estimated based on area under the curve of ten replicates determined by using growthcurver[4]. The box boundaries represent the first and third quartiles of the distribution and box-plot whiskers span 1.5 times the interquartile range of the distribution. Outliers are denoted as dots outside whiskers. Statistical significances were tested with a Wilcoxon two-sided test comparing the values of all strains against strain ANSES32139, which does not encode the $bla_{\text{OXA-181}}$ carbapenemase.

### Supplementary Tables

**Supplementary table 1** Characteristics of the *E. coli* isolates analysed in this work (Excel file)

**Supplementary table 2**. *E. coli* genome sequences retrieved from public databases (Excel file)

**Supplementary table 3**: Genomic features of the *E. coli* ST410 isolate *Ec*MAD

| | Replicon | Size (bp) | Number of CDS | Antibiotic resistance genes |
|---|---|---|---|---|
| Chromosome | | 4,747,851 | 4,452 | $bla_{CMY-2}$ |
| p*Ec*MAD1 | IncFIB, IncFIA, IncQ1 | 98,473 | 15 | *dfrA17, mph(A), aadA5, sul1, tet(B), aac(3)-IId, aac(6')Ib-cr,* $bla_{OXA-1}$, $bla_{CTX-M-15}$, $bla_{TEM-1B}$, *sul2, strA, strB* |
| p*Ec*MAD2 | IncX3, ΔColKP3 | 51,479 | 65 | *qnrS1*, $bla_{OXA-181}$ |
| p*Ec*MAD3 | Col (BS512) | 2,088 | 2 | - |

**Supplementary table 4** Antibiotics susceptibility testing of the ST410 isolate *Ec*MAD

| | |
|---|---|
| Resistant | Amoxicillin, Ticarcillin, Piperacillin, Cefepime, Cefalotin, Amoxicillin-Clavulanate, Ceftazidime, Piperacillin-Tazobactam, Cefoxitin, Cefuroxime, Ticarcillin-Clavulanate, Aztreonam, Moxalactam, Cefotaxime, Streptomycin, Clarythromycin, Tetracycline, Erythromycin, Rifampicin, Ciprofloxacin, Nalidixic Acid, Trimethoprim, Tobramycin |
| Intermediate | Mecillinam, Ertapenem, Kanamycin and Gentamicin |
| Susceptible | Imipenem, Meropenem, Doripenem, Amikacin, Azithromycin, Chloramphenicol, Tigecycline and Colistin |

**Supplementary table 5**. Distribution and effect of point mutations in the OXA-181 *E. coli* ST410 subclade MRCA

| Region | SNPs | Synonymous | Non-Synonymous | Start/Stop | Functional SNP* | IG§ |
|---|---|---|---|---|---|---|
| Non-recombinant | 97 | 35 | 48 | 2 | 25 | 12 |
| Recombinant& | 1648 (197) | 1324 (169) | 210 (16) | 2 | 9 (1) | 112 (11) |

*As determined by the SIFT algorithm[5], §Intergenic regions, &In bracket the number of SNPs in the 16.5 kb region of the *dcw* gene cluster common to the five recombinant regions in *Ec* ST410 isolates.

**Supplementary table 6**. Mutations predicted with a functional effect using the SIFT algorithm[5]

| Locus Tag | Gene | Product | Mutation | SIFT Score* | Median Info[&] | Chromosomal region | Functional class |
|---|---|---|---|---|---|---|---|
| EcMAD_00039 | *caiD* | crotonobetainyl-CoA hydratase | S181N | 0.01 | 2.75 | Non-recombinant | Nitrogen metabolism |
| EcMAD_00067 | *yabI* | hypothetical protein | G254V | 0 | 3.38 | Recombinant | Conserved hypothetical |
| EcMAD_00068 | *thiQ* | thiamin ABC transporter - ATP binding subunit | T66M | 0.03 | 2.76 | Recombinant | Transporter |
| EcMAD_00075 | *setA* | sugar / lactose efflux transporter SetA | V27G | 0.01 | 2.76 | Recombinant | Transporter |
| EcMAD_00088 | *ftsI* | essential cell division protein FtsI; penicillin-binding protein 3 | I536L | 0.02 | 2.75 | Recombinant | Cell envelope |
| EcMAD_00134 | *yadE* | putative polysaccharide deacetylase lipoprotein | P171S | 0.01 | 2.77 | Recombinant | Cell envelope |
| EcMAD_00143 | *htrE* | putative outer membrane usher protein | V662I | 0.05 | 2.75 | Recombinant | Cell envelope |
| EcMAD_00152 | *hrpB* | putative ATP-dependent helicase | C332S | 0.01 | 2.76 | Recombinant | DNA metabolism |
| EcMAD_00152 | *hrpB* | putative ATP-dependent helicase | P553L | 0.01 | 2.76 | Recombinant | DNA metabolism |
| EcMAD_00155 | *fhuC* | iron (III) hydroxamate ABC transporter - ATP binding subunit | T72A | 0.03 | 2.77 | Recombinant | Transporter |
| EcMAD_00362 | *brnQ* | branched chain amino acid transporter BrnQ | D385N | 0 | 2.77 | Non-recombinant | Transporter |
| EcMAD_00854 | *nfsA* | NADPH nitroreductase monomer | R203H | 0 | 2.77 | Non-recombinant | Stress response |
| EcMAD_00986 | *rarA* | acid phosphatase | A300V | 0 | 2.77 | Non-recombinant | DNA metabolism |
| EcMAD_01114 | *ymdA* | putative protein | I21S | 0.03 | 2.83 | Non-recombinant | Conserved hypothetical |
| EcMAD_01472 | *tehA* | TehA TDT transporter | P246L | 0 | 2.77 | Non-recombinant | Transporter |
| EcMAD_01480 | *hicB* | antitoxin of the HicA-HicB toxin-antitoxin system | T34A | 0 | 2.75 | Non-recombinant | Stress response |
| EcMAD_01535 | *dosP* | c-di-GMP phosphodiesterase, heme-regulated | G524D | 0.02 | 2.76 | Non-recombinant | Stress response |
| EcMAD_01541 | *yddB* | putative porin protein | G406C | 0 | 2.78 | Non-recombinant | Cell envelope |
| EcMAD_01796 | *astE* | succinylglutamate desuccinylase | A45V | 0.03 | 2.78 | Non-recombinant | Carbon metabolism |
| EcMAD_01900 | *edd* | phosphogluconate dehydratase | R312H | 0.05 | 3.22 | Non-recombinant | Carbon metabolism |
| EcMAD_02162 | *yegV* | putative kinase | I181T | 0 | 2.77 | Non-recombinant | Conserved hypothetical |
| EcMAD_02201 | *yohC* | putative inner membrane protein | L134Q | 0 | 2.78 | Non-recombinant | Conserved hypothetical |
| EcMAD_02264 | *ccmB* | protoheme IX ABC transporter - membrane subunit CcmB | G147S | 0 | 2.77 | Non-recombinant | Transporter |
| EcMAD_02280 | *ompC* | outer membrane porin C | R191L | 0.03 | 3.14 | Non-recombinant | Cell envelope |
| EcMAD_02311 | *glpB* | glycerol-3-phosphate dehydrogenase, membrane anchor subunit | V289M | 0.02 | 2.76 | Non-recombinant | Carbon metabolism |
| EcMAD_02425 | *emrK* | EmrKY-TolC multidrug efflux transport system - membrane protein | N248I | 0 | 2.75 | Non-recombinant | Transporter |
| EcMAD_02511 | *maeB* | malate dehydrogenase | A172S | 0.04 | 3.14 | Non-recombinant | Carbon metabolism |
| EcMAD_02714 | *nrdE* | ribonucleoside-diphosphate reductase 2 | F84L | 0.01 | 3.3 | Non-recombinant | DNA metabolism |
| EcMAD_02760 | *hycI* | hydrogenase 3 maturation protease | A63V | 0.02 | 2.79 | Non-recombinant | Carbon metabolism |
| EcMAD_02802 | *casC* | Cascade subunit C | P125S | 0 | 2.75 | Non-recombinant | DNA metabolism |
| EcMAD_03071 | *gss* | glutathionylspermidine amidase / glutathionylspermidine synthetase | P100S | 0.01 | 2.81 | Non-recombinant | Nitrogen metabolism |
| EcMAD_03159 | *ebgR* | EbgR DNA-binding transcriptional repressor | I98T | 0 | 2.77 | Non-recombinant | C. metabolism Regulation |
| EcMAD_03286 | *yrbG* | inner membrane protein YrbG | G219V | 0.01 | 2.76 | Non-recombinant | Conserved hypothetical |
| EcMAD_03840 | *emrD* | multidrug efflux transporter EmrD | G323D | 0 | 2.76 | Non-recombinant | Transporter |

* Ranges from 0 to 1, mutation is predicted to be functional for values equal or below 0.05. [&]Median for the whole alignment of the information content calculated for each position. This value ranges from 0, when all 20 amino acids are identified to 4.32 when only one amino acid is found.

**Supplementary table 7**: *ftsI, ompC* and *ompF* mutations in STs encompassing CP-*Ec* isolates (Excel file)

**Supplementary table 8**. Antibiotic resistance gene content of ST410 isolates analysed in Fig. 7*

| | *qnr*B4 | *qnr*S1 | *aac*(3)-IId | *aac*(6')Ib-cr | *aad*A5 | *bla*CMY-2 | *bla*CMY-42 | *bla*CTX-M-15 | *bla*DHA-1 | *bla*OXA-1 | *bla*OXA-181 | *bla*TEM-1B | *dfr*A17 | *mph*(A) | *strA* | *strB* | *Sul1* | *sul2* | *tet*(A) | *tet*(B) |
|---|---|---|---|---|---|---|---|---|---|---|---|---|---|---|---|---|---|---|---|---|
| 83B9 | + | + | + | + | + | + | | + | + | + | + | + | + | + | + | + | + | + | | + |
| *Ec*MAD | | + | + | + | + | + | | + | | + | + | + | + | + | + | + | + | + | | + |
| 94G8 | | + | | | + | | + | + | | | + | | + | + | | | + | + | + | |
| 92B7 | | + | | + | + | | | + | | + | + | | + | + | | | + | | + | |
| 93G1 | | + | | + | + | | | + | | + | + | | + | + | | | + | | + | |
| 32139 | | | + | + | + | | | + | | + | | + | + | + | + | + | + | + | | + |

*β-lactamase genes are highlighted with colours; + indicates that the gene is present.

**Supplementary table 9**. Oligonucleotides used in this work

| Gene | Name | Sequence (5'-3') | Experiment |
|---|---|---|---|
| *ompC* | ompC-F | GCGAAGGCATGACCAACAAC | qRT-PCR |
| *ompC* | ompC-R | AGGTCTGGGTGTACTGAGCA | qRT-PCR |
| *ompF* | ompF-F | ACCAACCGCAAGAAGCTCA | qRT-PCR |
| *ompF* | ompF-R | TTGGTGTAAGCGATGGACGG | qRT-PCR |
| *recA* | recA-F | GAAGACCGTTCCATGGATGT | qRT-PCR |
| *recA* | recA-R | GCGTGTTCAGCATCGATAAA | qRT-PCR |
| *ftsI* | Mut1ftsI | C*C*G*G*TCAGGGTTAATTTGCTGTAGCGTGCCACGTCTTTGATTTCGTGGCCGTTAATTCGATAGTTAATTCGATAAGGAATGGTATTGAGTA§ | TM-MAGE |
| *ftsI* | Mut2ftsI | C*C*A*T*GATGGCACCAAAGACCGGCGCGGAAACGGCGCCGCCGTAGTATTTACCCGCCTGCGGATCGTTGAGAACAACAACCAGCGCGAAGC§ | TM-MAGE |
| *ftsI* | Mut1ftsIwt_F | ACCCCGGTCAGGGTTAATTC | Mutant screening PCR |
| *ftsI* | Mut1ftsIM_F | ACCCCGGTCAGGGTTAATTT | Mutant screening PCR |
| *ftsI* | Mut1ftsI_R | ACCATCACCGACGTGTTTGA | Mutant screening PCR |
| *ftsI* | Mut2ftsIwt_F | CTTCGCGCTGGTTGTTGTTA | Mutant screening PCR |
| *ftsI* | Mut2ftsIM_F | CTTCGCGCTGGTTGTTGTTC | Mutant screening PCR |
| *ftsI* | Mut2ftsI_R | CCTGTCCCCTCGCCTTGATT | Mutant screening PCR |
| *ftsI* | Mut1ftsIwt2_F | CGTGGCCGTTAATTCGATAA | Mutant screening PCR |
| *ftsI* | Mut1ftsIM2_F | CGTGGCCGTTAATTCGATAG | Mutant screening PCR |
| *ftsI* | Mut1ftsI2_R | CGAAAGAGGCGATGCGTAAC | Mutant screening PCR |
| *ompC* | ompC_Xba_F | *CAGAATCTAGA*GTTTTGACATTCAGTGCTGTCA | Cloning in pACY184 |
| *ompC* | ompC_Hind_R | *AACATAAGCTT*TCATGCGAACGGTCGCAAGA | Cloning in pACY184 |
| *ompC* | ompC_F2 | CAGACATTCAGAAATGAATGACGGT | Sequence verification |
| *ompC* | ompC_R2 | AGTGCCACGCGGGTAAAGCT | Sequence verification |
| *ompC* | ompC_St38_F | TAGTGCTCATGGCGAAGGTA | Sequence verification |
| *ompC* | ompC_St38_R | GTACCGATCAGGCCAGTGT | Sequence verification |
| *pACY184* | pACY184_F | AAGAGATTACGCGCAGACCA | Sequence verification |
| *pACY184* | pACY184_R | ATAGTGACTGGCGATGCTGT | Sequence verification |

§ * indicates a phosphorothioate bonds

**Supplementary table 10**: Mutations detected in MG1655 strains derivatives mutated in *ftsI*.

| *ftsI* mutation | Mutant name | Position | Mutation | Gene affected | *Effect |
|---|---|---|---|---|---|
| Control§ | Magect_1 | 1,64,569 | A=>G | *ydfA* | NS |
| | | 1,842,545 | C=>T | *gdhA* | NS |
| | | 2,236,418 | G=>A | *preA* | NS |
| | | 2,426,060 | A=>G | *hisJ* | S |
| YRIN | Mage_1a | 92,411 | Ins 12nt | *ftsI* | **YRIN** |
| | | 777,276 | $(G)_{6\rightarrow7}$ | *tolA* | FS |
| | | 979,184 | C=>T | *mukB* | S |
| | | 4,261,075 | G=>T | *yjbM* | NS |
| E349K | Mage_2b | 92,457 | G=>A | *ftsI* | **E349K** |
| | | 380,022 | $(G)_{10\rightarrow9}$ | *frmR* | IG |
| | | 1,475,884 | A=>G | *ydbD* | S |
| I532L | Mage_3c | 93,006 | A=>C | *ftsI* | **I532L** |
| YRIN+E349K | Mage_4b | 92,411 | Ins 12nt | *ftsI* | **YRIN** |
| | | 92,457 | G=>A | *ftsI* | **E349K** |
| E349K + I532L | Mage_5a | 92,457 | G=>A | *ftsI* | **E349K** |
| | | 93,006 | A=>C | *ftsI* | **I532L** |
| | | 21,878 | T=>C | *ribF* | NS |
| | | 2,958,079 | A=>G | *ptrA* | NS |
| YRIN+E349K + I532L | Mage_6b | 92,411 | Ins 12nt | *ftsI* | **YRIN** |
| | | 92,457 | G=>A | *ftsI* | **E349K** |
| | | 93,006 | A=>C | *ftsI* | **I532L** |
| | | 994,214 | C=>A | *ssuC* | NS |

§Control strain was obtained by submitting the MGF strain to TM-MAGE steps but without incorporating mutations in *ftsI*. §NS, non-synonymous, S, synonymous; FS Frameshift, IG intergenic. Mutations resulting of site directed mutagenesis are indicated in red

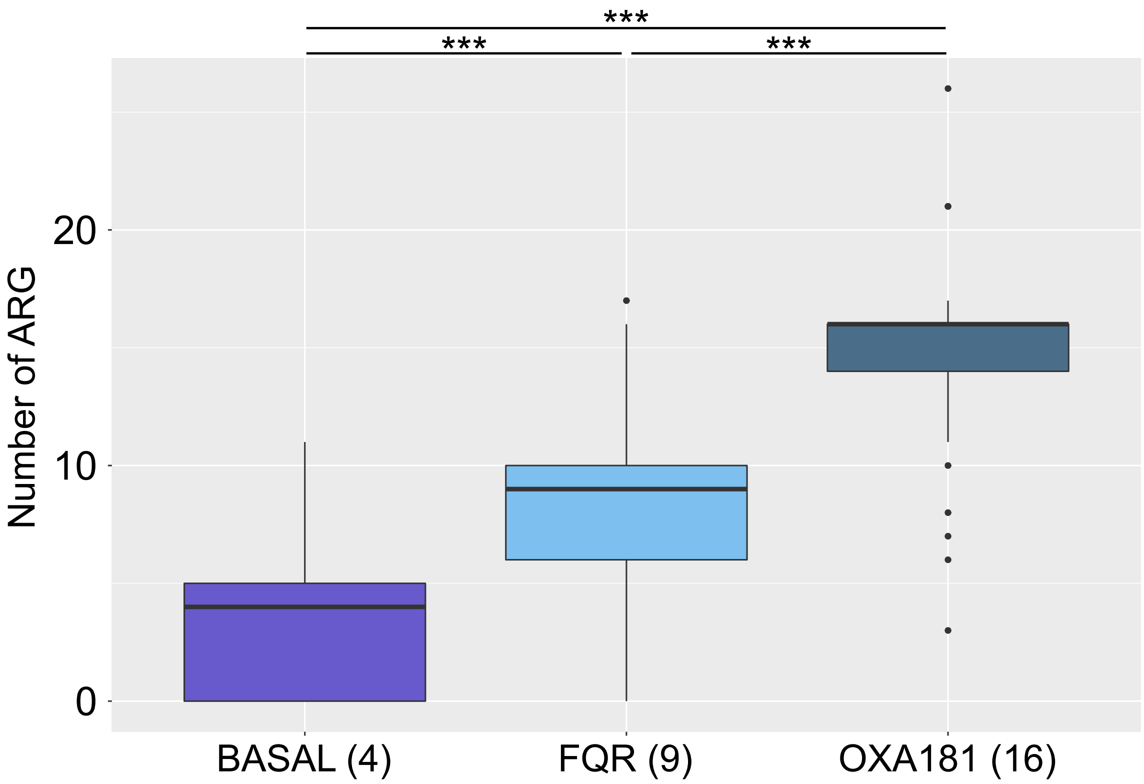


Fig. S1

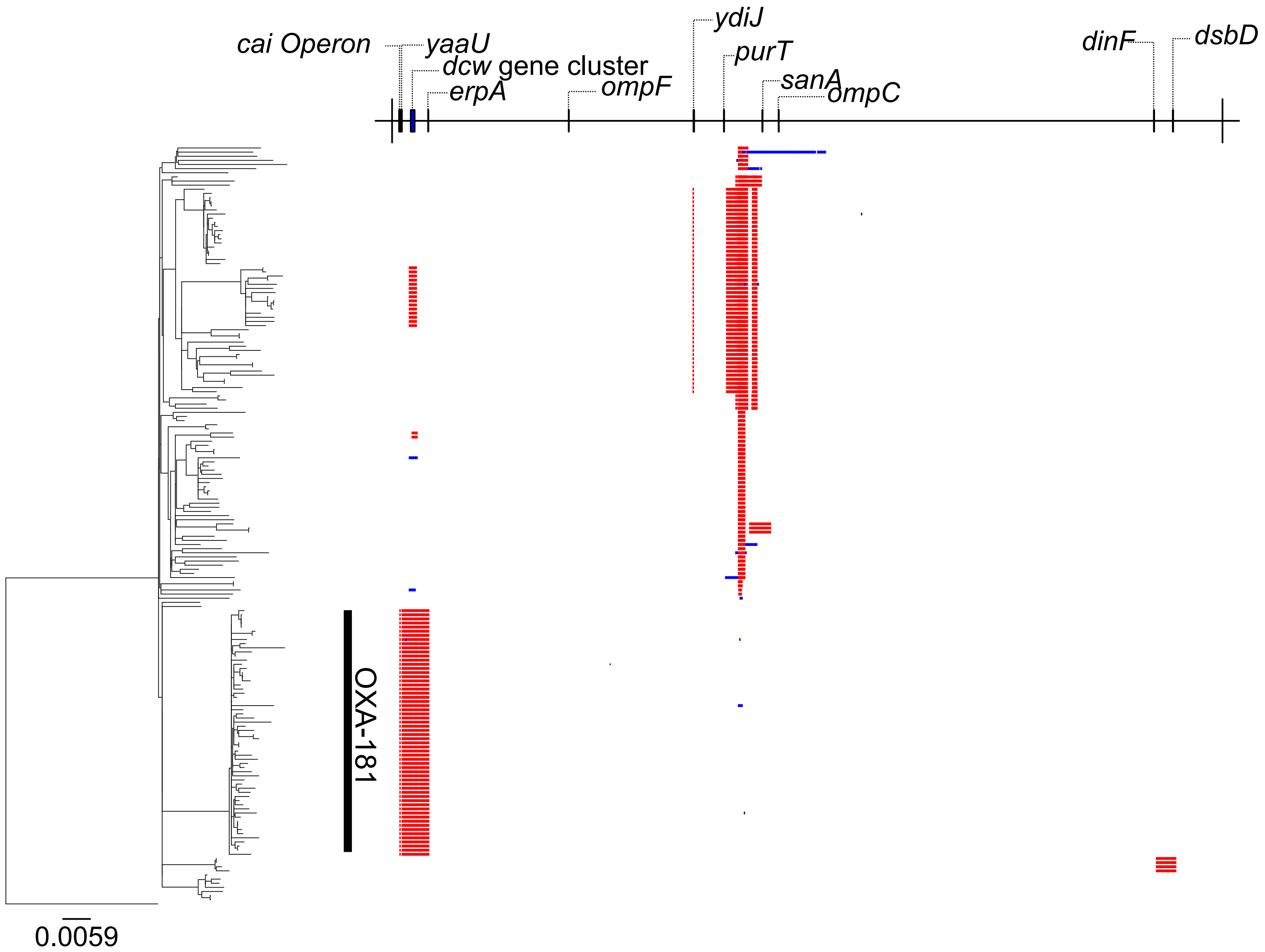
cai Operon
yaaU
dcw gene cluster
erpA
ompF
ydiJ
purT
sanA
ompC
dinF
dsbD
OXA-181
0.0059

Fig. S2

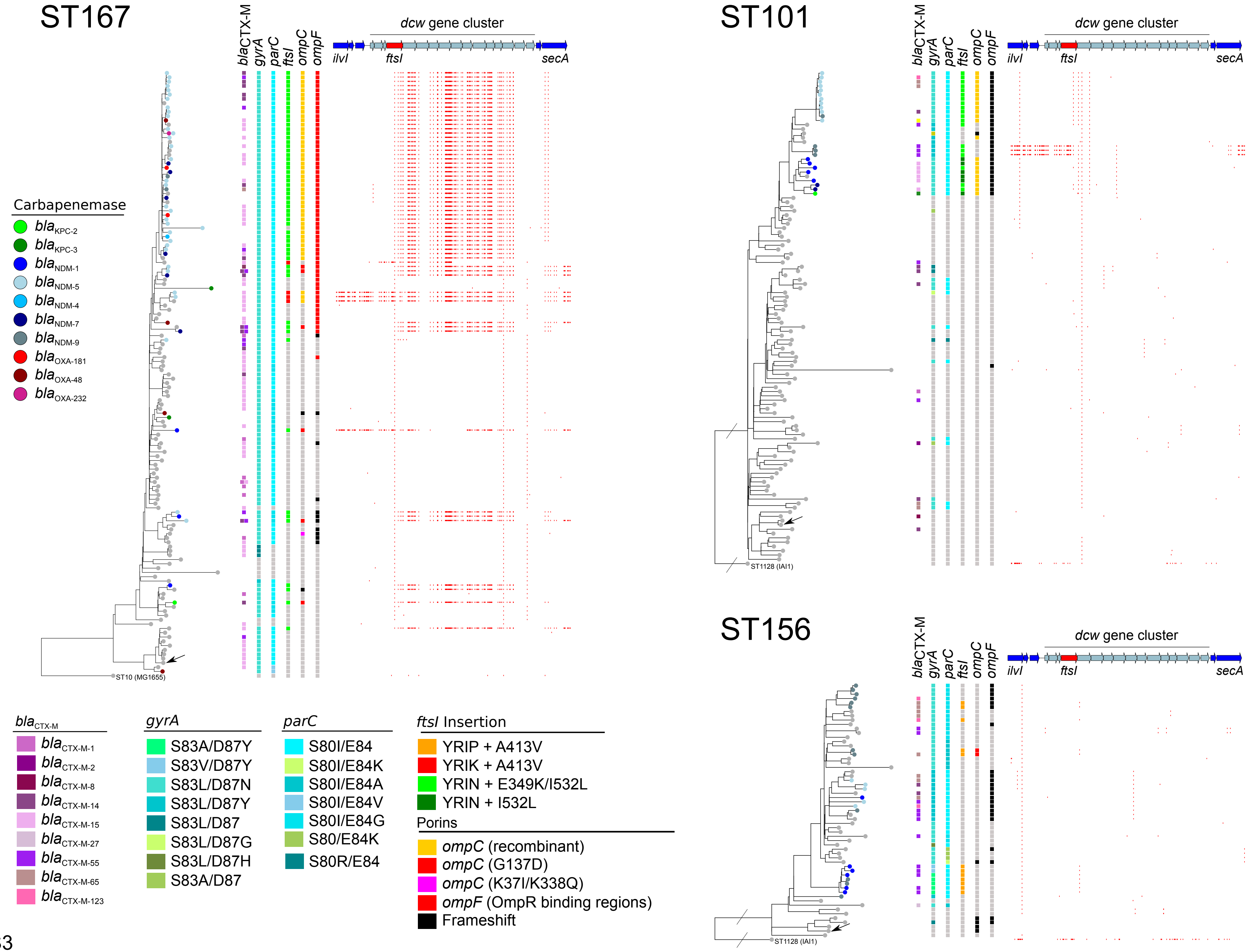

ST167
ST101
ST156
blaCTX-M
gyrA
parC
ftsI
ompC
ompF
dcw gene cluster
ilvI
ftsI
secA
ST10 (MG1655)
ST1128 (IAI1)
Carbapenemase
blaKPC-2
blaKPC-3
blaNDM-1
blaNDM-5
blaNDM-4
blaNDM-7
blaNDM-9
blaOXA-181
blaOXA-48
blaOXA-232
blaCTX-M
blaCTX-M-1
blaCTX-M-2
blaCTX-M-8
blaCTX-M-14
blaCTX-M-15
blaCTX-M-27
blaCTX-M-55
blaCTX-M-65
blaCTX-M-123
gyrA
S83A/D87Y
S83V/D87Y
S83L/D87N
S83L/D87Y
S83L/D87
S83L/D87G
S83L/D87H
S83A/D87
parC
S80I/E84
S80I/E84K
S80I/E84A
S80I/E84V
S80I/E84G
S80/E84K
S80R/E84
ftsI Insertion
YRIP + A413V
YRIK + A413V
YRIN + E349K/I532L
YRIN + I532L
Porins
ompC (recombinant)
ompC (G137D)
ompC (K37I/K338Q)
ompF (OmpR binding regions)
Frameshift


Fig. S3

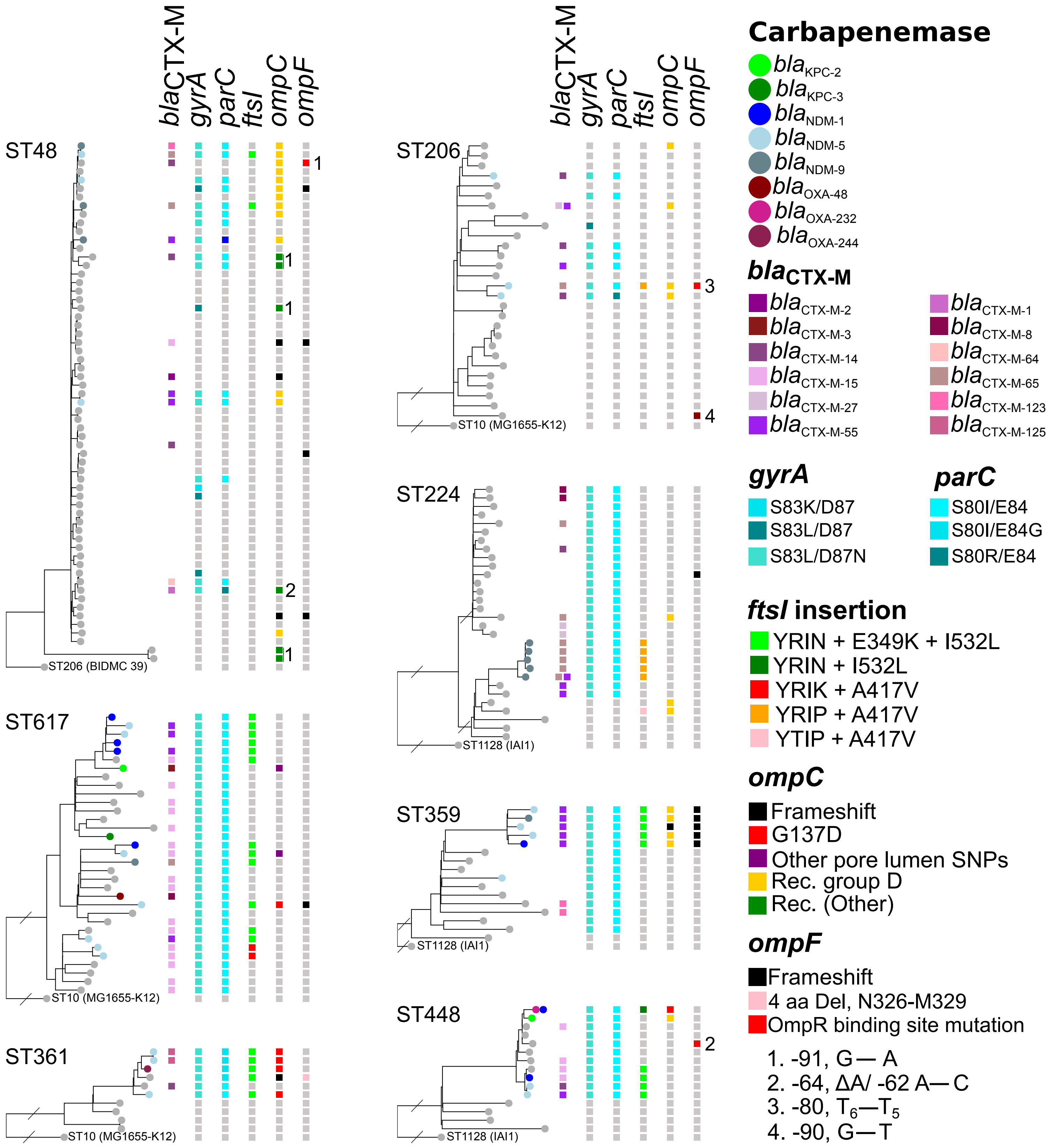
ST48
ST206 (BIDMC 39)
ST617
ST10 (MG1655-K12)
ST361
ST10 (MG1655-K12)
ST206
ST10 (MG1655-K12)
ST224
ST1128 (IAI1)
ST359
ST1128 (IAI1)
ST448
ST1128 (IAI1)
blaCTX-M
gyrA
parC
ftsI
ompC
ompF
Carbapenemase
blaKPC-2
blaKPC-3
blaNDM-1
blaNDM-5
blaNDM-9
blaOXA-48
blaOXA-232
blaOXA-244
blaCTX-M
blaCTX-M-2
blaCTX-M-3
blaCTX-M-14
blaCTX-M-15
blaCTX-M-27
blaCTX-M-55
blaCTX-M-1
blaCTX-M-8
blaCTX-M-64
blaCTX-M-65
blaCTX-M-123
blaCTX-M-125
gyrA
S83K/D87
S83L/D87
S83L/D87N
parC
S80I/E84
S80I/E84G
S80R/E84
ftsI insertion
YRIN + E349K + I532L
YRIN + I532L
YRIK + A417V
YRIP + A417V
YTIP + A417V
ompC
Frameshift
G137D
Other pore lumen SNPs
Rec. group D
Rec. (Other)
ompF
Frameshift
4 aa Del, N326-M329
OmpR binding site mutation
1. -91, G— A
2. -64, ΔA/ -62 A— C
3. -80, T6—T5
4. -90, G—T

Fig. S4

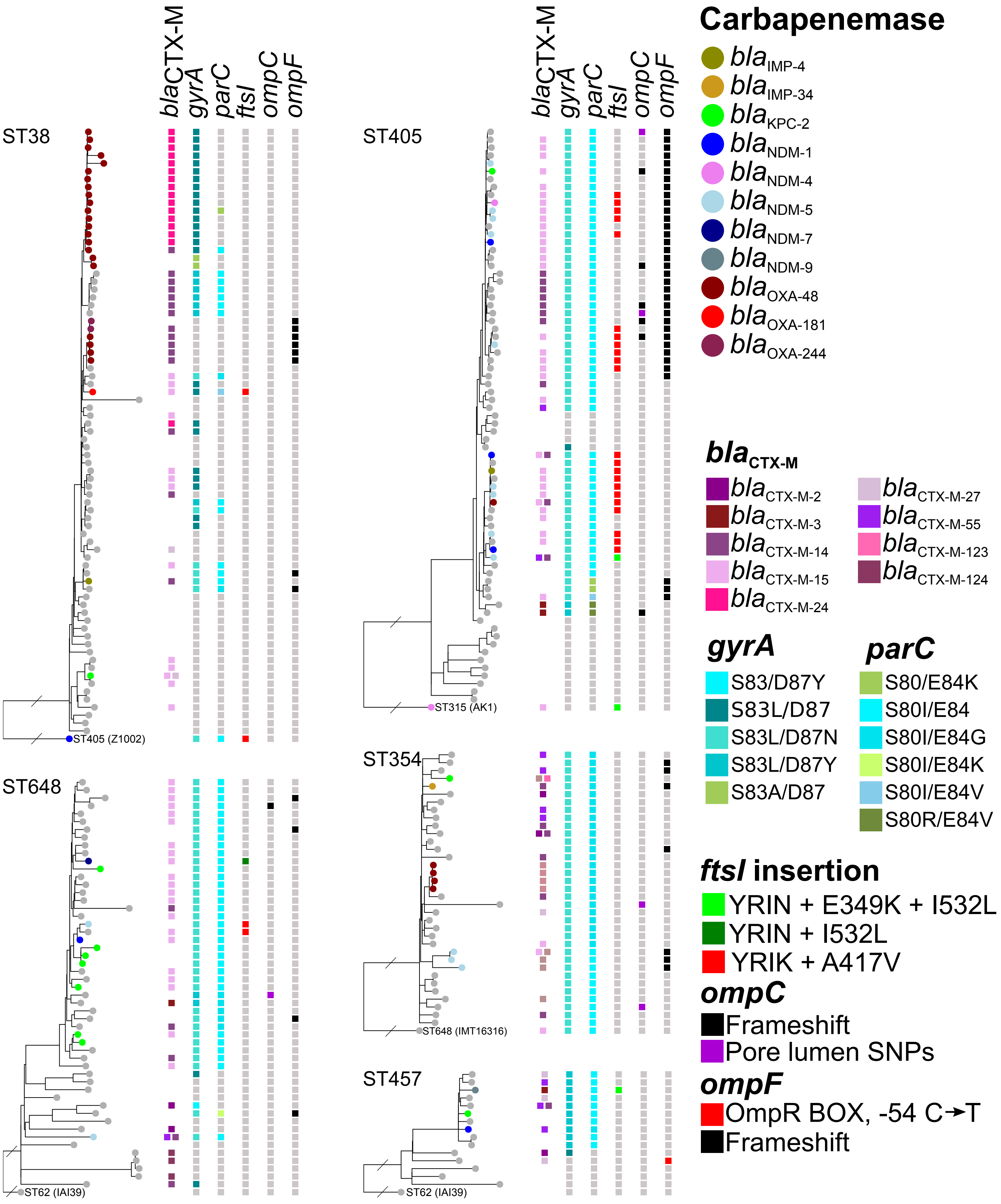
blaCTX-M
gyrA
parC
ftsI
ompC
ompF
ST38
ST405 (Z1002)
ST405
ST315 (AK1)
ST648
ST62 (IAI39)
ST354
ST648 (IMT16316)
ST457
ST62 (IAI39)
Carbapenemase
blaIMP-4
blaIMP-34
blaKPC-2
blaNDM-1
blaNDM-4
blaNDM-5
blaNDM-7
blaNDM-9
blaOXA-48
blaOXA-181
blaOXA-244
blaCTX-M
blaCTX-M-2
blaCTX-M-3
blaCTX-M-14
blaCTX-M-15
blaCTX-M-24
blaCTX-M-27
blaCTX-M-55
blaCTX-M-123
blaCTX-M-124
gyrA
S83/D87Y
S83L/D87
S83L/D87N
S83L/D87Y
S83A/D87
parC
S80/E84K
S80I/E84
S80I/E84G
S80I/E84K
S80I/E84V
S80R/E84V
ftsI insertion
YRIN + E349K + I532L
YRIN + I532L
YRIK + A417V
ompC
Frameshift
Pore lumen SNPs
ompF
OmpR BOX, -54 C→T
Frameshift

Fig. S5

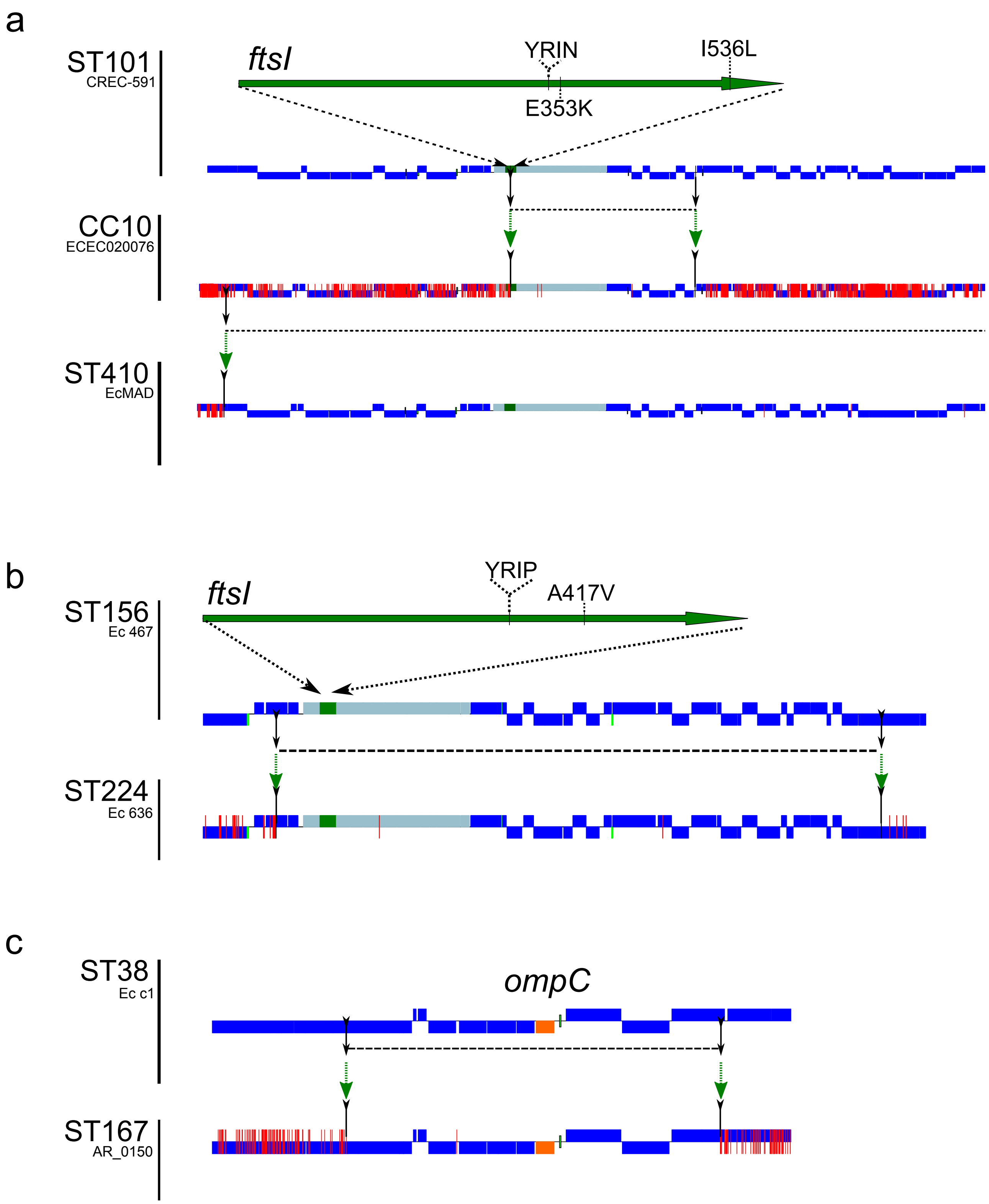
a
ST101
CREC-591
ftsI
YRIN
I536L
E353K
CC10
ECEC020076
ST410
EcMAD
b
ST156
Ec 467
ftsI
YRIP
A417V
ST224
Ec 636
c
ST38
Ec c1
ompC
ST167
AR_0150

Fig. S6

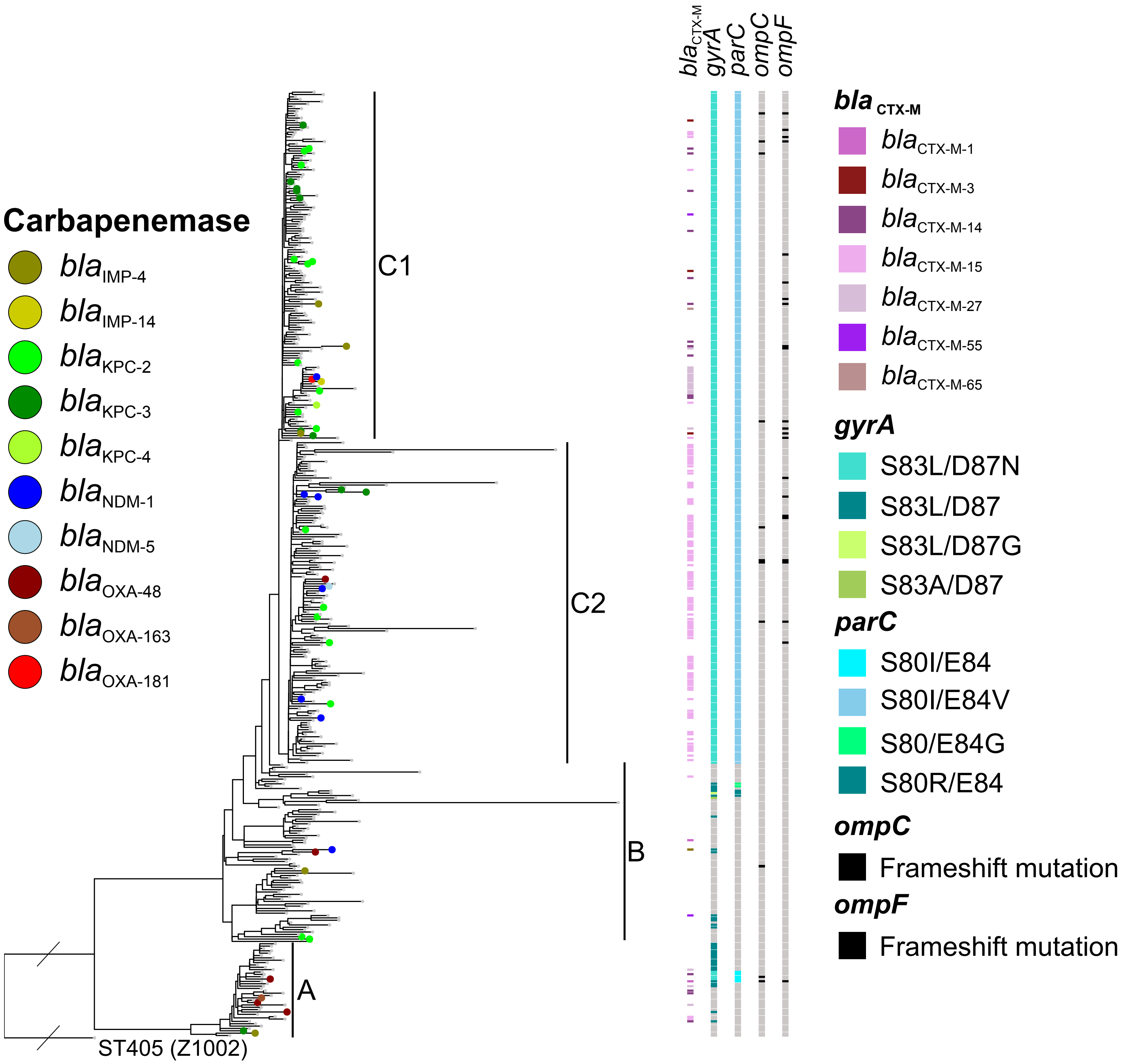
Carbapenemase
blaIMP-4
blaIMP-14
blaKPC-2
blaKPC-3
blaKPC-4
blaNDM-1
blaNDM-5
blaOXA-48
blaOXA-163
blaOXA-181
C1
C2
B
A
ST405 (Z1002)
blaCTX-M
gyrA
parC
ompC
ompF
bla CTX-M
blaCTX-M-1
blaCTX-M-3
blaCTX-M-14
blaCTX-M-15
blaCTX-M-27
blaCTX-M-55
blaCTX-M-65
gyrA
S83L/D87N
S83L/D87
S83L/D87G
S83A/D87
parC
S80I/E84
S80I/E84V
S80/E84G
S80R/E84
ompC
Frameshift mutation
ompF
Frameshift mutation

Fig. S7

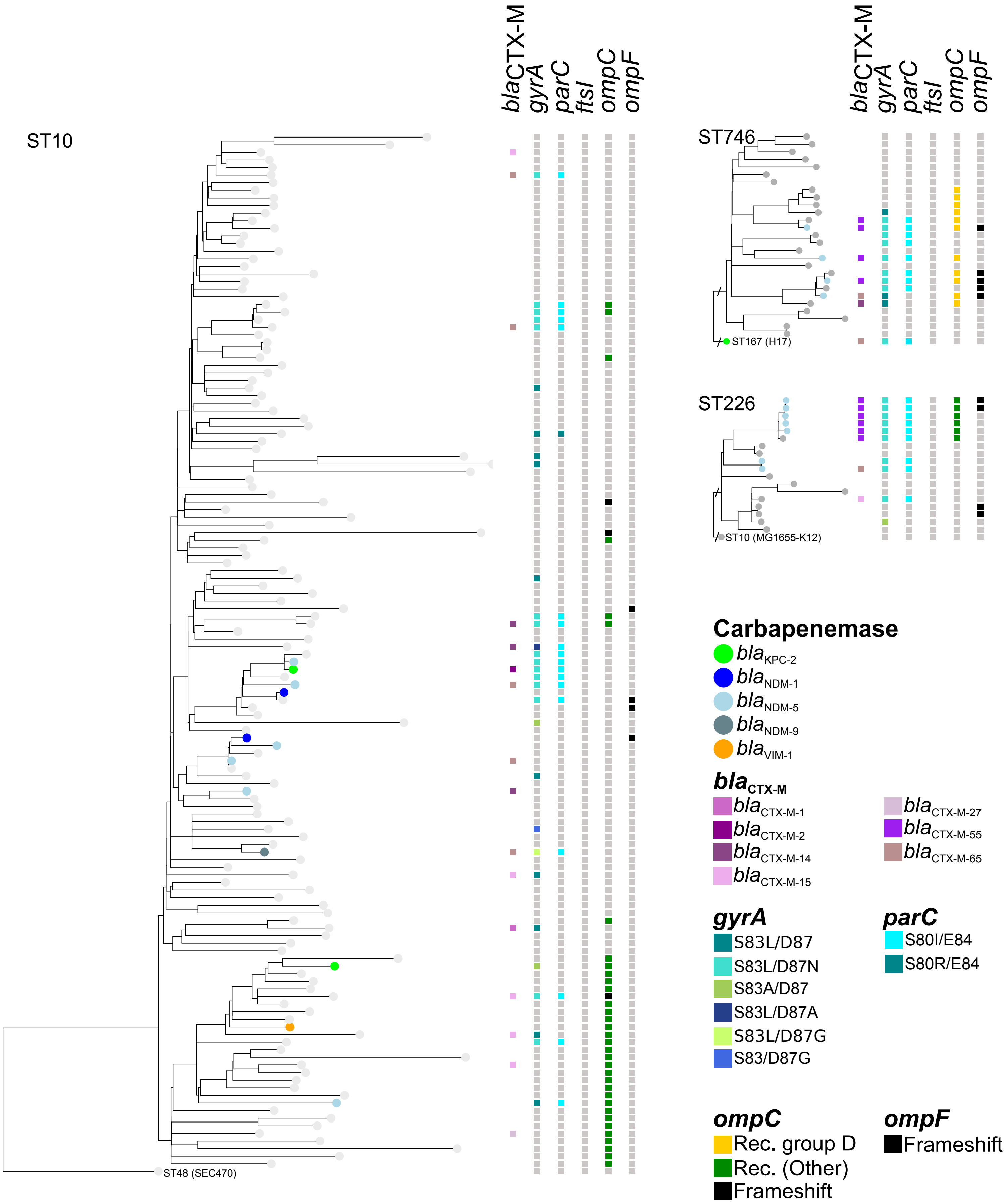

ST10
ST48 (SEC470)
blaCTX-M gyrA parC ftsI ompC ompF
ST746
ST167 (H17)
ST226
ST10 (MG1655-K12)
Carbapenemase
blaKPC-2
blaNDM-1
blaNDM-5
blaNDM-9
blaVIM-1
blaCTX-M
blaCTX-M-1
blaCTX-M-2
blaCTX-M-14
blaCTX-M-15
blaCTX-M-27
blaCTX-M-55
blaCTX-M-65
gyrA
S83L/D87
S83L/D87N
S83A/D87
S83L/D87A
S83L/D87G
S83/D87G
parC
S80I/E84
S80R/E84
ompC
Rec. group D
Rec. (Other)
Frameshift
ompF
Frameshift


Fig. S8

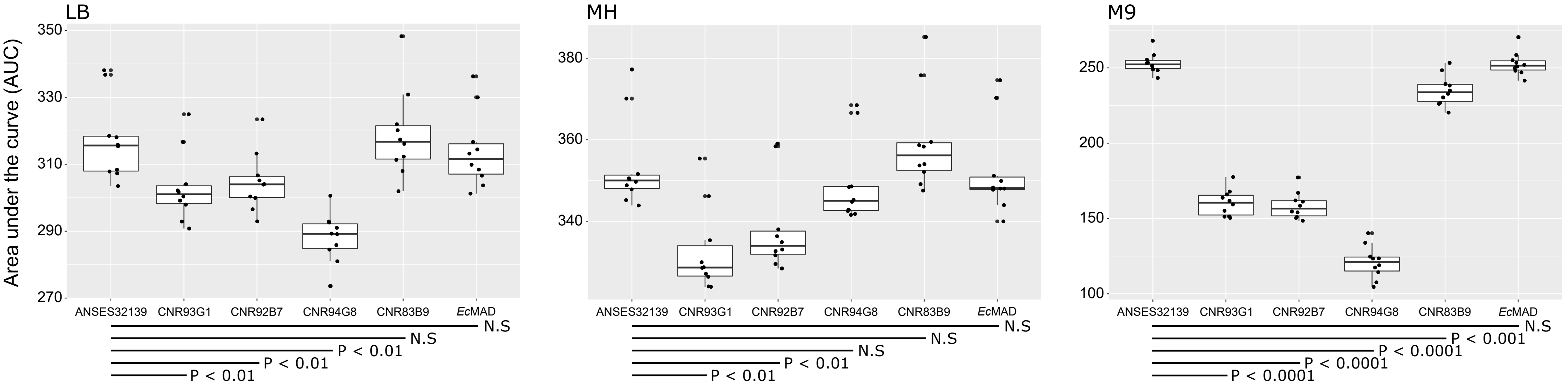
LB
MH
M9
Area under the curve (AUC)
350
330
310
290
270
380
360
340
250
200
150
100
ANSES32139
CNR93G1
CNR92B7
CNR94G8
CNR83B9
EcMAD
N.S
N.S
P < 0.01
P < 0.01
P < 0.01
N.S
N.S
N.S
P < 0.01
P < 0.01
N.S
P < 0.001
P < 0.0001
P < 0.0001
P < 0.0001

Fig. S9
